# Red blood cell-derived extracellular vesicles mediate intercellular communication in ischemic heart failure

**DOI:** 10.1101/624841

**Authors:** Avash Das, Nedyalka Valkov, Ane M. Salvador, Ivan Kur, Olivia Ziegler, Ashish Yeri, Fernando Camacho Garcia, Shulin Lu, Aushee Khamesra, Chunyang Xiao, Rodosthenis Rodosthenous, Guoping Li, Srimeenakshi Srinivasan, Vasilis Toxavidis, John Tigges, Louise C. Laurent, Stefan Momma, Ionita Ghiran, Saumya Das

**Author notes:** These authors contributed equally (Ionita Ghiran, Saumya Das). Correspondence to: 1. Saumya Das, Simches 3, Cardiovascular Research Center, Massachusetts General Hospital, 185 Cambridge Street, Boston, MA 02114., 2. Ionita Ghiran, Center for Life Sciences, Beth Israel Deaconess Medical Center, 3 Blackfan Circle, Boston, MA 02215.

## Abstract

Extracellular vesicles (EV) mediate intercellular signaling by transferring their cargo to recipient cells. Red blood cell (RBC)-derived EVs constitute a significant proportion of circulating EVs and have been implicated in regulating immune responses. Here, we describe a transgenic mouse model for fluorescent-based mapping of RBC-EV target cells based on the functional transfer of EV-contained Cre-recombinase to target cells. In a murine model of ischemic heart failure, we detect an increase in RBC-EV-targeted cardiomyocytes in the hearts and microglial cells in the brains. Cells targeted by RBC-EVs present an enrichment of genes implicated in cell proliferation and metabolism pathways compared to non-recombined (non-targeted) cells. Cardiomyocytes targeted by RBC-EVs are more likely to demonstrate cellular markers of DNA synthesis and proliferation, suggesting functional significance of EV-mediated signaling. In conclusion, we leverage our mouse model for mapping of RBC-EV targets in murine ischemic heart failure to demonstrate quantitative and qualitative changes in RBC-EV recipients.

## Introduction

Extracellular vesicles (EVs) are cell-derived membranous structures (100-1,000 nanometer in diameter) comprising exosomes and microvesicles (Maas et al., 2017, van Niel et al., 2018, Tkach and Thery, 2016). EVs carry diverse cargo including lipids, proteins, long non-coding RNA (lncRNA) and microRNA (miRNA) (Hong et al., 2009; Mathivanan et al., 2012; Valadi et al., 2007) that can be transferred to recipient cells (Pironti et al., 2015, Takasaki et al., 1977, Ridder et al., 2014) to mediate intercellular communication. Several recent studies have demonstrated the importance of small RNAs contained within EVs as important functional mediators of intercellular signaling. In this regard, miRNAs, generally known as negative regulators of gene expression inside the cell (Bartel, 2009) have been shown to constitute a significant proportion of RNA present in EVs (Bellingham et al., 2012). Recent studies have demonstrated the transfer of EV-miRNAs to recipient cells with subsequent alterations in target mRNA expression and phenotype of recipient cells (Bang et al., 2014, Jae et al., 2015, Ying et al., 2017).

Most of our understanding about EV function comes from studies using EVs derived either from cell culture conditioned media or biological fluids, and their subsequent administration in animal models to assess functional changes. This is very unlikely to reflect their natural composition and endogenous function *in vivo*. Our knowledge regarding EV target cells, and the functional consequences for the target cells *in vivo* remains largely unknown, mainly due to the lack of suitable tools and techniques to track EV targets *in vivo*. We have previously shown that functional Cre mRNA can be packaged in exosomes released by Cre recombinase-expressing cells and transferred to EV-recipient reporter cells, subsequently mediating Cre-dependent recombination to allow expression of a reporter (Ridder et al., 2014). This system has allowed for mapping of targets of hematopoietic cell-derived EVs at baseline and in a model of peripheral inflammation.

RBCs are a significant source of EVs in plasma, with release of EVs triggered by membrane complement activation and calcium influx, providing a mechanism for delivery of EVs at the site of inflammation (Kuo et al., 2017). This mechanism of EV release can be leveraged to generate large quantities of pure RBC-EVs ex-vivo for experimental use.

In addition, RBC-EVs express unique membrane proteins, such as blood group antigens and glycophorins, which allows for antibody-based sorting of RBC-specific EVs and profiling of their cargo. These properties make RBC-EVs attractive for mechanistic studies (Kuo et al., 2017). RBC-EVs play an immunomodulatory role with a demonstrated function in regulating monocyte-mediated activation of endothelial cells (Straat et al., 2016b), modulating T-cell-monocyte immune synapse (Danesh et al., 2014), and altering endothelial cells via transfer of functional microRNAs in the context of malaria infection (Mantel et al., 2016).

Adverse cardiac remodeling following myocardial infarction (MI) can eventually lead to heart failure (Fishman et al., 2010; WRITING GROUP MEMBERS et al., 2010). Cardiac remodeling comprises complex molecular, cellular and interstitial changes (Cohn et al., 2000) that are mediated by multiple signaling pathways such as apoptosis, inflammation, and fibrosis (Honold and Nahrendorf, 2018, Travers et al., 2016) and interactions between multiple cell types, although the complex pathophysiology of cardiac remodeling remains elusive (Azevedo et al., 2016). Recent therapeutic strategies that target inflammation to reduce heart failure (Ridker et al., 2017) suggest that interaction between hematopoietic cells and cardiac cells may be important in cardiac remodeling. Notably, recent evidence demonstrates the prognostic significance of RBC-EVs in post-MI patients (Cheow et al., 2016, Giannopoulos et al., 2014, Giannopoulos et al., 2017), and plasma miRNAs derived from circulating hematopoietic cells (HCs), have been implicated in cardiac remodeling in several studies (Collino et al., 2010, Wen et al., 2012). However, the functional contribution of RBC-EVs as mediators of *intercellular signalin*g (Thery et al., 2009) that contribute to post-MI LV remodeling remains unexplored, largely due to the limitations in tools for in-vivo tracking of EVs (Gangadaran et al., 2017).

Here, we report the application of a fluorescent-based EV-target mapping technique based on the Cre-LoxP system (Ridder et al., 2014) to track RBC-EVs in a transgenic murine model. The EpoR-Cre transgenic mouse (*Cre* expression under the erythropoietin receptor promoter (Kerenyi et al., 2013, Heinrich et al., 2004)), when crossed with the Rosa26 mTomato/mGFP (Muzumdar et al., 2007) mouse, leads to mGFP expression in RBC, erythropoietic progenitor cells and platelets to some degree (as they arise from megakaryocyte-erythrocyte precursors, MEPs) (Klimchenko et al., 2009). The RBCs in turn, secrete mGFP-positive EVs that also contain functional Cre protein. Transfer of functional Cre to target cells allows for identification of RBC-EV target cells in vivo. We leverage this EV-mapping model to study the targets of RBC-EVs at baseline and in a murine ischemic heart failure model (following ischemia/reperfusion/infarction or I/R). We show the qualitative and quantitative alteration in RBC-EVs targets with I/R and demonstrate possible remote functional consequences of RBC-EV targeting. Taken together, our study is the first to show the distribution and target cell types of endogenous RBC-EVs *in vivo* and can be used by investigators to study the functional consequences of RBC-EV mediated signaling.

## Results

### Murine model for fluorescent-based mapping of RBC-EVs target cells

To study EV-mediated communication between RBCs and different tissues, we crossed erythroid lineage-specific *Cre* knock-in mice (EpoR-Cre) (Heinrich et al., 2004) with membrane-targeted tandem dimer (td) Tomato/membrane-targeted GFP (mT/mG) mice (Muzumdar et al., 2007) to generate double transgenic mice (EpoR-Cre/mTmG). In the absence of Cre, mTmG mice express only Tomato on the cell membranes of all cells, but after exposure to Cre Recombinase (i.e., in the EpoR-Cre/mTmG mice), tdTomato (mTd) is excised, resulting in loss of tdTomato and gain of mGFP expression in Cre-expressing cells (**Figure 1A**). As previously described (Ridder et al., 2014), EVs from the Cre-expressing cells transfer functional Cre to EV-target cells, resulting in Cre-mediated recombination and expression of mGFP in the target cells. Furthermore, EVs from the Cre-expressing cells (in this case RBC-EVs) can be identified as GFP-positive EVs by fluorescence microscopy or nano-flow cytometry (Danielson et al., 2016). Accordingly, fluorescence microscopy of the peripheral blood smear and cell free-plasma (CFP) from EpoR-Cre/mTmG double transgenic mice confirmed that all of the RBCs in the circulation and a substantial fraction of the EVs in plasma were mGFP+, as expected. (**Fig. 1B)**. Flow cytometry analysis of isolated blood cells (following removal of the buffy coat from peripheral blood) of EpoR-Cre/mTmG mice revealed the presence of Ter 119 (circulating erythroid-specific marker), confirming the presence of mGFP^+^ RBCs in the peripheral circulation (**Figure S1A**). Ter 119 immuno-negative but GFP^+^ cells are contaminating platelets (see also Fig. 1B) that arise from the megakaryocyte-erythrocyte precursor (MEPs, known to express EpoR, hence are GFP^+^) (Klimchenko et al., 2009). Nanoflow cytometry of the plasma from EpoR-Cre/mTmG mice plasma confirmed the presence of mGFP^+^ EVs, while plasma from mTmG mice showed only mTd^+^ EVs suggesting that GFP^+^-EVs are mainly of RBC origin (**Figure 1C**) with some contribution from platelets. Expectedly, no fluorescence was detected in the EpoR-Cre mice (**Figure S1B**). Quantification of EVs based on their fluorescence from EpoR-Cre/mTmG mice (mTd^+^, mGFP^+^ and mTd^+^/mGFP^+^) is represented in **Figure S1C** and showed significant individual variation (ranging from 19.28% to 64.86%) of the circulating GFP^+^ EVs.

**Figure 1.**
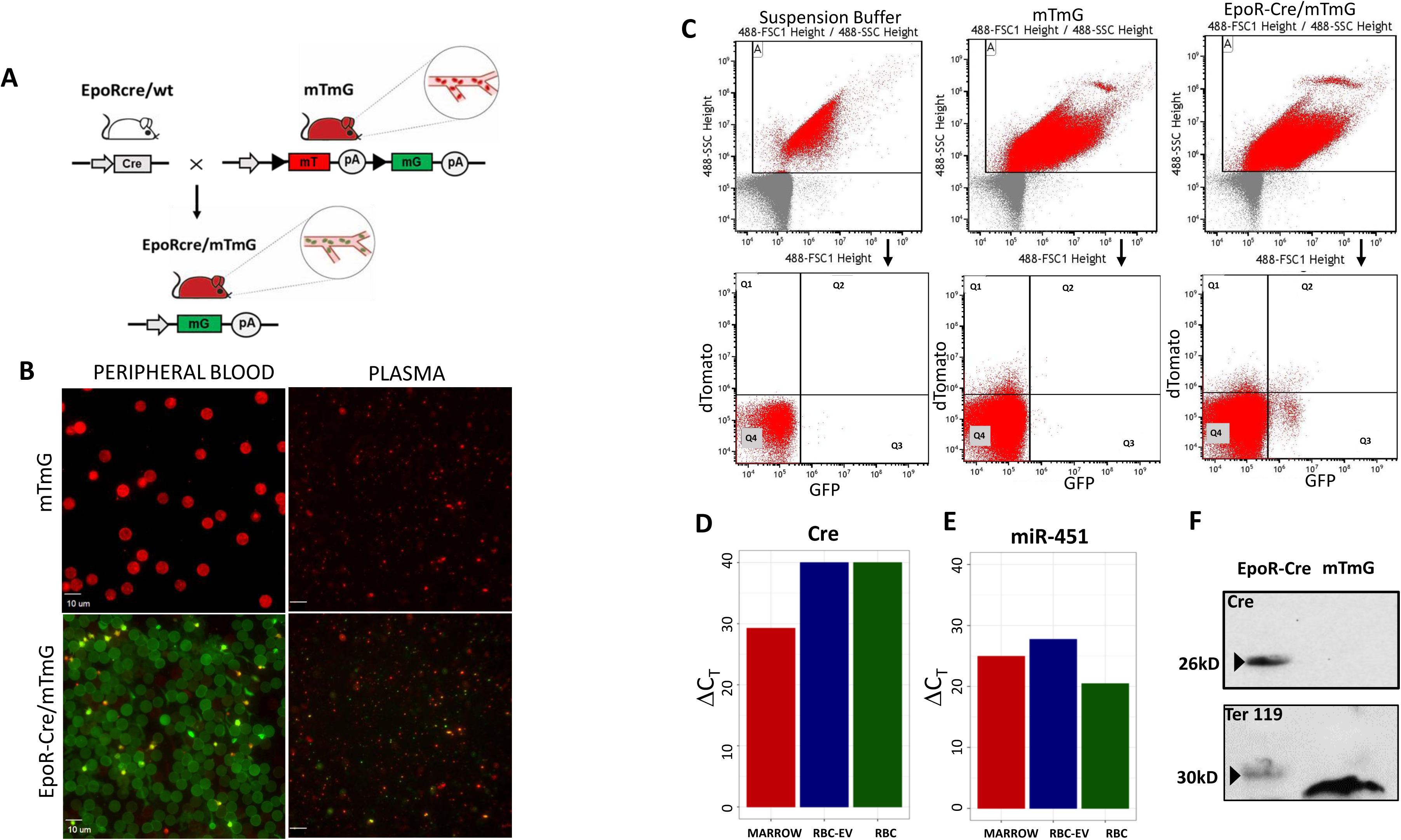
Baseline characterization of RBC-EVs in transgenic murine model. (A) Schematic representation of the experimental murine models. Erythroid lineage specific conditional *Cre* knock-in mice (EpoR-Cre mice, BALB/c background) were crossed with membrane-targeted Tomato/membrane-targeted GFP (mTmG^Rosa26^ mice, C57BL6/Sv129 background) to produce double transgenic (EpoR-Cre/mTmG) off-springs. Activated *Cre* mediates the removal of membrane-targeted tandem dimer Tomato (mT) sequence allowing the expression of membrane-targeted green fluorescent protein (mG) in RBCs. (B) High magnification fluorescence images (63X) of fresh prepared peripheral blood smear and CFP smear from mTmG and EpoR-Cre/mTmG mice to characterize the mature circulating RBCs, and RBC-EVs (n=4 in each group). Small bright GFP positive dots in the peripheral blood smear represent platelets. Scale bar: 10 µm. (C) Nanoflow cytometric characterization of CFP from mTmG mice and EpoR-Cre/mTmG mice, showing presence of mGFP^+^ RBC-EVs in EpoR-Cre/mTmG mouse plasma. The auto fluorescence from the Suspension Buffer (Phosphate Buffer Saline) has been accounted for, as background. Shown are Td on the y-axis and GFP in the x-axis and divided in four quadrants [Q1 is mTd^+^ (red); Q2 is mTd^+^/mGFP^+^ (double positive); Q3 is mGFP^+^ (green) and Q4 is background. (D) Gene expression levels of Cre-recombinase mRNA obtained from bone marrow, RBC-EVs and RBC from EpoR-Cre mice. Expression is shown as raw ΔCT (ΔCT of 29.25 for bone marrow, >40 for RBC-EVs and >40 for RBCs) over biological duplicates. (E) Gene expression levels of micro-RNA (miR)-451a (documented to be abundantly expressed in matured RBCs) in bone marrow, RBC-EVs and RBC from EpoR-Cre mice. Data are shown as raw ΔC_T_ over biological duplicates (ΔC_T_ of 24.99 for bone marrow, 27.72 for RBC-EVs and 20.45 for RBCs respectively). (F) Western blot of RBC-EVs protein lysates from EpoR-Cre and mTmG mice probed with the indicated antibodies: Cre-recombinase and Ter-119 (mature RBC-specific marker). The exposure time to detect Cre-recombinase was 200 secs and Ter-119 was 100 secs. Data shown are representative of 3 replicated independent experiments.

To demonstrate the presence of Cre-recombinase in RBC-EVs, we assessed Cre mRNA and protein expression levels in RBC-EVs derived from EpoR-Cre and mTmG mice. Real time PCR showed the presence of Cre-recombinase mRNA in bone marrow cells, but their absence in peripheral blood and RBC-EVs (**Figure 1D, E**), consistent with the general absence of mRNAs in RBCs (Moras et al., 2017) but the presence of the prototypical RBC miRNA miR-451 in all three compartments (bone marrow cells, RBCs and RBC-EVs). However, western blot confirmed the presence of Cre-recombinase protein in RBC-EVs (**Figure 1F)**, suggesting that Cre protein present in RBCs can be incorporated into RBC-EVs.

Next, we demonstrated the transfer of functional Cre*-*recombinase protein via RBC-EVs occurs *in vitro*. We treated either primary adult dermal fibroblast derived from mTmG mice (Seluanov et al., 2010) (all mTd^+^ at the time of isolation) or a Cre-reporter HEK cell line we generated with total EVs isolated from EpoR-Cre or mTmG mouse CFP plasma through serial density gradient ultracentrifugation. We observed mGFP^+^ dermal fibroblasts or HEK cells (consistent with Cre-dependent recombination) in the EpoR-Cre EV treatment group, but no recombination events in the mTmG EV treated group, demonstrating RBC-EV mediated functional transfer of Cre (**Figure S1D).**

### Mapping the targets of RBC-EVs

EVs have recently been demonstrated to mediate intercellular communication (Ridder et al., 2015, Ratajczak et al., 2006, Valadi et al., 2007). To determine the cellular targets of RBC-EVs in our EV-mapping mouse model, we leveraged existing methodology on successful administration of EVs systemically or locally (Alexander et al., 2015; Ohno et al., 2013; Ridder et al., 2014; Wiklander et al., 2015). Intraperitoneal (IP) injection of mGFP^+^ RBC-EVs, generated ex-vivo from isolated and purified RBCs from EpoR-Cre/mTmG mice using complement activation, into wild type C57BL/6 mice resulted in the presence of mGFP^+^ EVs in the circulation 48 hours following i.p. administration, as demonstrated by fluorescence microscopy from cell free plasma (**Figure S2A**). Next, complement-generated RBC-EVs from EpoR-Cre mice (expected to contain functional Cre protein) or from mTmG ‘control’ mice were transfused into the mTmG reporter mice. Tissue and plasma from the recipient mice were analyzed 7 days after transfusion (**Figure 2A**). Flow cytometry of the blood and nano flow cytometry of cell-free plasma (**Figure 2B, C**) confirmed the presence of mGFP^+^ RBCs in the circulation and mGFP^+^ RBC-EVs in the plasma of the mTmG mice treated with EpoR-Cre RBC-EVs but not in control mTmG RBC-EVs (which were not expected to contain Cre). Together, these results demonstrated that i.p. delivery of RBC-EVs form EpoR-Cre mice result in recombination and expression of mGFP in both progenitor and mature RBCs and subsequent generation of mGFP^+^ EVs in the plasma, thereby phenocopying the EpoR-Cre/mTmG double transgenic mice.

**Figure 2.**
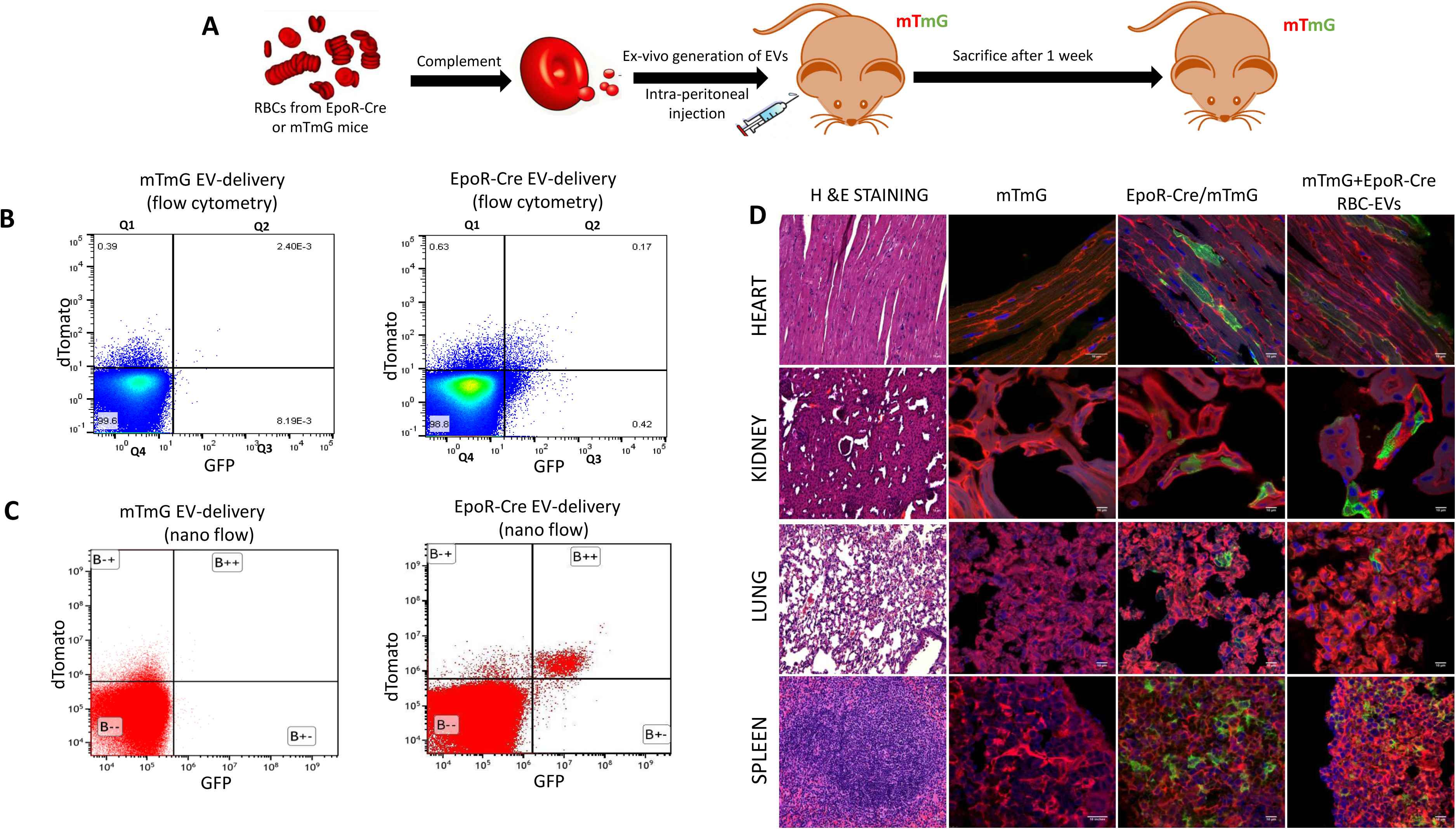
*In-vivo* transfer of functional Cre-recombinase through RBC-EVs. (A) Schematic representation of experimental design for intraperitoneal injection in mTmG mice with complement-generated RBC-EVs. (B) Flow cytometry scatter-plots of peripheral blood from mTmG mice transfused with mTmG RBC-EVs and EpoR-Cre RBC-EVs. Shown are Td on the y-axis and GFP in the x-axis and divided in four quadrants [Q4 is background, Q1 is mTd (red), Q3 is mGFP+ (green) and Q2 is mTd+/mGFP+ (double-positive)]. (C) Nanoflow cytometric characterization of CFP from mTmG mice transfused with EpoR-Cre RBC-EVs. Shown are Td on the y-axis and GFP in the x-axis and divided in four quadrants [B-- is background, B-+ is mTd+(red), B+- is mGFP+ (green) and B++ is mTd+/mGFP+ (double-positive)]. (D) Low magnification fluorescence microscopy (20X) of paraffin embedded Hematoxylin and Eosin stained sections of heart, kidney, lung and spleen from mTmG, EpoR-Cre/mTmG and mTmG mice transfused with EpoR-Cre RBC-EVs to characterize organ-specific tissue architecture. Corresponding, high magnification (63X) confocal microscopy of fixed frozen section of heart, kidney, lung and spleen from mTmG, EpoR-Cre/mTmG and mTmG mice transfused with EpoR-Cre RBC-EVs to map their fluorescence profile [mTomato (red) and mGFP (green)] at baseline and after transfusion. The findings were separately verified using biological replicates (n=4 for mTmG, n=6 for EpoR-Cre/mTmG and n=4 for mTmG mice transfused with EpoR-Cre RBC-EVs).

To determine cell-types and organs targeted by RBC-EVs, we analyzed sections of other organs harvested from the EpoR-Cre/mTmG double transgenic mice. Cre-mediated recombination (mTd^+^ to mGFP^+^ transformation) was noted in cells in the heart, kidney, lungs, spleen and pericytes in brain, suggesting transfer of RBC-EV molecular cargo (including Cre) at baseline (**Figure 2D, Figure S2B**), while interestingly, the liver showed no recombination (**Figure S2B**). To exclude the possibility of ‘leaky’ (Jaisser, 2000) or ectopic expression of Cre under the Erythropoietin (Epo) promoter leading to recombination in GFP^+^ cells (Ott et al., 2015, Sanchis-Gomar et al., 2014, Sinclair et al., 2010), or cell-fusion between circulating blood cells and the recombined cells (Yang et al., 2012, Zhang et al., 2004) we also analyzed the relevant organs harvested from the mTmG mice transfused with EpoR-Cre RBC-EVs, as described above. These transfusion experiments also allowed us to probe the targets of RBC-EVs specifically (as opposed to the double transgenic EpoR-Cre/mTmG mice that also had circulating platelet EVs). The heart, kidney, lungs and spleen showed the same profile of recombined cells as the EpoR-Cre/mTmG mice (**Figure 2D, Figure S2B)**, while the liver remained without any GFP-positive cells, confirming that recombination in these organs is caused by functional Cre transfer through RBC-EVs specifically, and is not mediated via EV-independent mechanisms. EV quantification by Nanoflow cytometry demonstrated the relative abundance of RBC-EVs compared with EVs from all other cell-types in the peripheral plasma of the transfused mTmG mice (**Figure S2C**): approximately 22.5% of EVs in the circulation were mGFP^+^, and hence arose from mGFP^+^ RBCs that were derived from erythrocyte progenitor cells or MEPs that had been targeted by the CRE-containing EVs that had been transfused. A quantitative count profile of the mTd^+^/mGFP^-^(red) and mTd^-^/mGFP^+^(green) cells at baseline in EpoR-Cre/mTmG mice compared with EpoR-Cre RBC-EV transfusion in mTmG mice for these organs are shown in **Figure S3** (40 high power field images for each organ per mice).

### RBC-EVs mediated signaling in post-infarct cardiac remodeling

RBCs have previously been implicated in vascular pathology such as atherosclerosis (Blum, 2009), thrombosis (Noh et al., 2010), and inflammation (Buesing et al., 2011). In addition, EVs derived from RBCs have been shown to increase under pathological stress (Ankarklev et al., 2014, Giannopoulos et al., 2014). Since there was EV-mediated cross-talk among RBC and cardiomyocytes under homeostatic milieu, and RBC-EVs are generated at the site of complement activation (as would happen following coronary thrombosis and myocardial infarction (Earis et al., 1985, Jordan et al., 2001, Rossen et al., 1988))), we hypothesized that RBC-EVs may play a role in post-infarction cardiac remodeling. We employed the well-validated model of ischemia/reperfusion/infarction (I/R, **Figure 3A**) to induce MI in EpoR-Cre/mTmG mice (Danielson et al., 2018, Melman et al., 2015), and assessed RBC-EV contribution to the injured heart 4 weeks after I/R.

**Figure 3.**
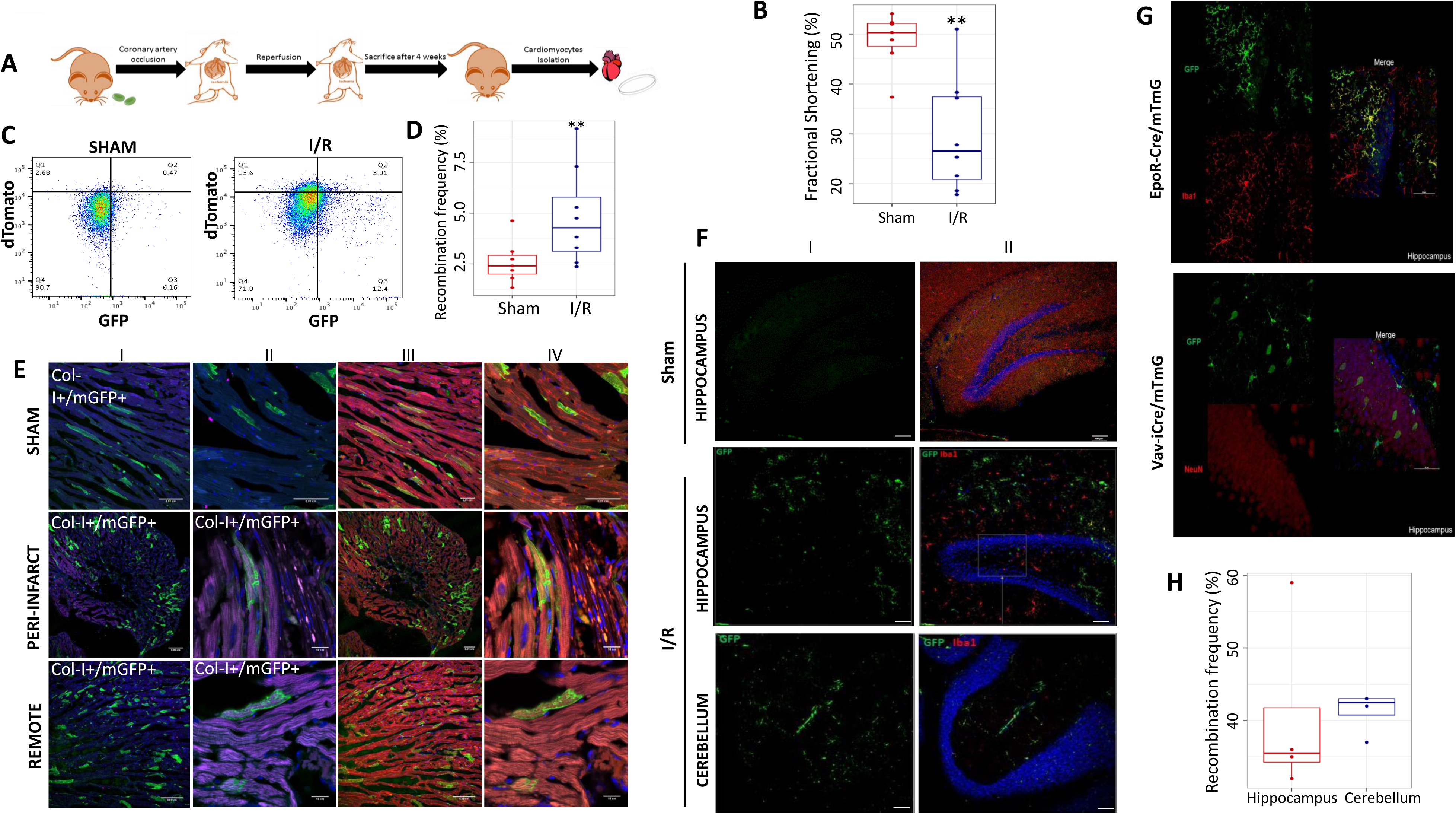
RBC-EV mediated intercellular communication in ischemic heart failure. (A) Schematic representation of experimental design for EpoR-Cre/mTmG mice undergoing I/R. (B) Box and whisker plots quantifying Fractional Shortening in Sham and I/R group of EpoR-Cre/mTmG mice (26.57±1.97% vs 50.3±1.97%, p=0.015) (n=8 for I/R and n=7 for Sham). Data are presented as Median with the interquartile range. (C) Representative flow cytometry scatter-plots of) CMs (viable, DAPI) from Sham and I/R EpoR-Cre/mTmG mice comparing the mGFP+ CMs between both groups. Shown are Td on the y-axis and GFP in the x-axis and divided in four quadrants [Q4 is background, Q1 is mTd+ (red), Q3 is mGFP+ (green) and Q2 is mTd+/mGFP+(double-positive)]. (n=7 for Sham and n=8 for I/R). (D) Box and whisker plots quantifying the recombination frequency of CMs in Sham and I/R group of EpoR-Cre/mTmG mice (2.41±0.37% vs 4.29±0.79%, p=0.013) n=8 for I/R and n=7 for Sham. Data are presented as Median with the interquartile range. (E) Representative fluorescent images of frozen ventricular heart sections of EpoR-Cre/mTmG I/Rmice from sham, remote and peri-infarct areas. (I) Low magnification (20X) and (II) corresponding high magnification (63X) images of heart sections stained with Collagen I (stains for fibrosis, pseudo-color Magenta) and DAPI staining of cell nucleus (blue). (III) Low magnification (20X) and (IV) corresponding high magnification (63X) images of heart sections stained for mTomato (red) and mGFP (green). Scale bar: 0.01 cm and 10 cm for low and high magnification images, respectively. (F) High magnification (63X) confocal microscopy along with 3D reconstruction and 2D projection of images from fixed paraffin-embedded brain tissue sections form EpoR-Cre/mTmG mice undergoing I/R. (I) Mapping of nuclei (DAPI, stained blue) and GFP+ cells (green) in hippocampus and cerebellum of EpoR-Cre/mTmG mice undergoing I/R (II) Immunostaining of the sections with Iba1 antibody (pseudo-color red, co-staining for resident microglia) and corresponding overlay of PE (red) channel to demonstrate Iba1+/mGFP+ cells (n = 4). Scale bar: 0.01 cm and 10 cm. (G) Immunostaining of brain tissue sections from EpoR-Cre/mTmG and vav-iCre mice undergoing I/R with Iba1 and Anti-NeuN antibody.).Corresponding high magnification (63X) confocal microscopy along with 3D reconstruction and 2D projection of images showing GFP+ cells in the hippocampus in the two groups (n=4). (H) Box and whisker plot quantifying Iba1+/mGFP+ microglia in the hippocampus and cerebellum of EpoR-Cre/mTmG mice undergoing I/R (42±0.02% vs 36±5; n=4). Data is presented as median with the interquartile range.

Echocardiographic measurements (Figure 3B) showed that I/R induced ventricular dysfunction in the mice as indicated by decrease in FS in the I/R group compared to sham. Nanoflow cytometry of the CFP did not show any significant difference in the count profile of plasma total EVs and RBC-EVs (mGFP^+^ EVs) between the I/R and sham groups mice at the 4-week time point (**Figure S4A, B, C**). Cardiomyocytes were isolated from the hearts of these animals as previously described (Ackers-Johnson et al., 2016). Flow cytometric characterization of viable cardiomyocytes showed a significant increase in the number of mGFP^+^ cardiomyocytes in the I/R group compared to their sham counterparts (**Figure 3C, D**). We next assessed spatial predisposition of the mGFP^+^ cardiomyocytes in the I/R group in respect to the infarcted area. At the 4 week-stage in the remodeling process, there was no distinct grouping/or increased density of the recombined GFP^+^ cardiomyocytes close to or around the infarct or peri-infarct area (as identified by collagen-immuno-reactive areas to highlight fibrotic areas). Recombined cardiomyocytes were seen even in the remote areas of the myocardium, suggesting that at this later time-point in remodeling, RBC-EVs targeted cardiomyocytes both in the peri-infarct and remote zones (**Figure 3E**). Microscopic count profile of the kidney, lung, and spleen did not show any significant difference between I/R, sham and baseline groups (**Figure S4D**).

To confirm whether the observed pattern of communication between RBCs and cardiomyocytes after I/R is mediated by RBC-EVs and not a result of cell-fusion (Yang et al., 2012; Zhang et al., 2004), we performed I/R on mTmG mice with injection of RBC-EVs during reperfusion (**Figure S5A**). The mice were sacrificed after 4 weeks and the viable cardiomyocytes isolated and characterized with Flow Cytometry and microscopy to demonstrate the presence of mGFP^+^ cardiomyocytes (**Figure S5B, C**).

### RBC-EV-mediated communication with brain after I/R

Our previous study had demonstrated communication between hematopoietic cells and brain cells (including neurons and microglia) in murine models of peripheral inflammation and peripheral arterial disease (Ridder et al., 2014) Given recent attention on the association between ischemic heart disease and neuro-cognitive disorders (Almeida et al., 2012, Chen et al., 2017, Lichtman et al., 2008, van der Wall and van Gilst, 2013), we were interested in exploring the role of RBC-EVs as mediators of communication between the periphery and the brain in the context of ischemic heart disease. Images of brains from EpoR-Cre/mTmG mice demonstrated the presence of mGFP^+/^Ib1a^+^ microglia in the cerebellum and hippocampus, 4 weeks after I/R (**Figure 3F)**. As a comparison we did not detect recombined microglia in the sham animals. Interestingly, the target of the RBC-EVs is different from the targets of other hematopoietic cells: recombination is seen in neurons as well as microglia in vav-iCre/ mTmG mice following peripheral inflammation (Ridder et al., 2014) or I/R (**Fig. 3G**), suggesting that the cellular origin of EVs may in part determine their target cells. The mGFP^+^/Ib1a^+^ microglia were more predominant in the cerebellum in comparison to hippocampus in EpoR-Cre/mTmG mice (**Figure 3H**). These findings demonstrate that cardiac injury induces a marked increase in targeting of microglia by RBC-EVs with possible differences between the hippocampus and the cerebellum.

### Functional effects of RBC-EV mediated signaling *in vitro*

Previous studies have demonstrated the importance of EV cargo RNAs, notably miRNAs in modulating recipient cell function (Valadi et al., 2007, Melo et al., 2014). We therefore sought to determine the small RNA content of RBC-EVs and whether it was altered in the context of myocardial infarction. RBC-EVs could be isolated by flow sorting based on the presence of the Glycophorin-A (CD235a) antigen on their surface; however, given the very low yield of RNA from flow sorting of a subset of EVs in plasma, this was not technically feasible in mice models. We explored RBC-EV content in a clinically well-characterized bio-repository of human plasma from patients with myocardial infarction, heart failure and supraventricular tachycardia. Small RNA-seq was used to analyze RNA content in Glycophorin-A (CD235a)-immunopositive EVs from the plasma of patients with type I myocardial infarction, chronic heart failure or control patients referred for supraventricular tachycardia (see clinical characterization of the patients in **Table S1**).

Due to the very low RNA yield leading low input reads, we analyzed the pooled RNAseq data for those samples that passed minimal QC (see methods) and identified a total of 105 distinct miRNAs in the patient plasma derived RBC-EVs, with no significant differences noted between the three cohorts grouped together. Among them, however, only a few miRNAs were highly expressed in all three cohorts. MiRNA-451a was the most enriched, comprising 11.7 % of the total miRNA reads in terms of reads per million (RPM) mapped to the transcriptome, followed by miR486-5p (9.4%) and let-7a-5p (2.70%) (**Figure 4A**). The targets of these top 20 miRNAs were identified using TargetScan v 7.1 and all target mRNAs with a cumulative context score less than 0.2 were included for Gene Ontology analysis using DAVID (Huang da et al., 2009). We identified enriched pathways involved in cell cycle, cell division, cell migration and proliferation (**Figure 4B**) suggesting that while RNA content of RBC-EVs does not appear to change in the context of cardiovascular disease, the transfer of the miRNA contents may regulate the pathways described above in recipient cells.

**Figure 4.**
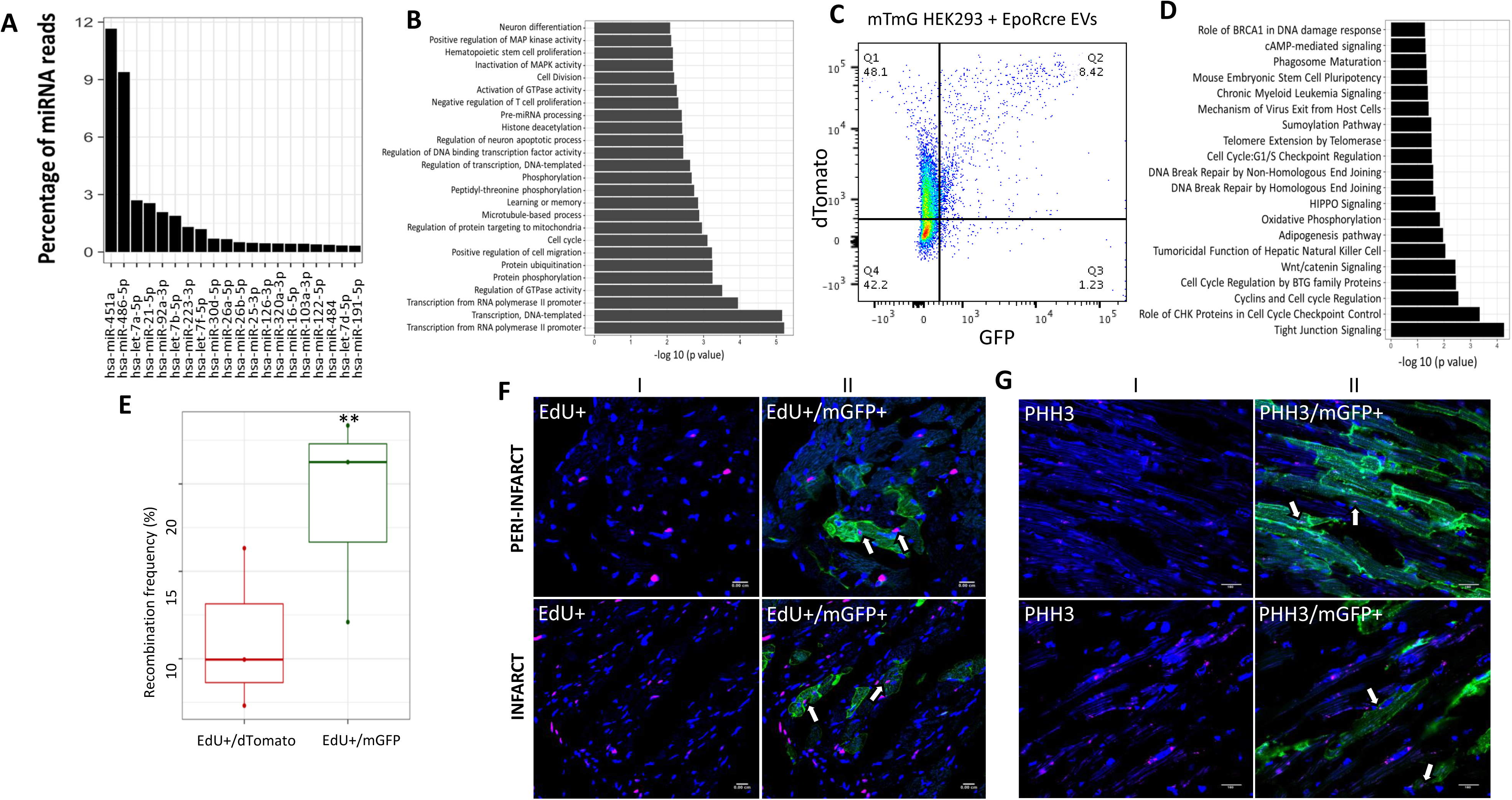
RBC-EVs modulate post-infarct cardiac remodeling. (A) Profiling of human miRNAs isolated from patient plasma derived RBC-EVs. Percentage of reads measured as reads per million (RPM) mapped to the transcriptome (RPM) for the top 20 miRNAs. hsa-miR-451a is the most enriched miRNA that accounts for ∼11.7% of the reads. (B) Bar plot representing the p-values for the top 20 Gene Ontology (GO) pathways enriched by targets of the top 20 expressing miRNAs in patient plasma derived RBC-EVs. Target genes were identified using TargetScan v7.1 and DAVID pathway analysis was used to evaluate the significance of the each GO pathway. (C) Flow cytometry scatter-plots of mTmG HEK293 cells treated with EpoR-Cre RBC-EVs for 36 hours. Shown are Td on the y-axis and GFP in the x-axis and divided in four quadrants [Q4 is background: Q1 is mTd+(red), Q3 is mGFP+ (green) and Q2 is mTd+/mGFP+ (double-positive)]; n= 3 independent replicates for each condition. (D) Ingenuity Pathway Analysis of enriched pathways for transcriptomic data for flow-sorted mGFP+ HEK293 cells and mTd+ HEK293 cells after treatment of mT/mG HEK293 cells with EpoR-Cre EVs and mTmG EVs for 36 hrs. (n=3 each). A total of 143 genes that were expressed with at least 25 RPM and a 2 fold change between mGFP+ HEK293 and mTd+ HEK293 cells were included for pathway analysis. (E) Box and whisker plot comparing EdU+/mTd+ and EdU+/mGFP+ CMs in the EpoR-Cre/mTmG mice undergoing I/R (21.25±2.82% vs 9.95±2.18%, p=0.056). n=3. Data are presented as median with the interquartile range. (F) Representative fluorescent images of frozen ventricular heart sections of Edu injected EpoR-Cre/mTmG mice with I/R from infarct and peri-infarct areas. (I) High magnification (63X) images showing EdU – positive cell nuclei (Alexa 647, pseudo-color Magenta) and nuclei stained with DAPI (blue) highlighting their overlap. (II) corresponding overlay of GFP channel demonstrating EdU+/mGFP+ CMs. n = 4 for I/R and n=3 for sham. (G) Representative fluorescent images of frozen ventricular heart sections ofEpoR-Cre/mTmG mice with I/R from infarct and peri-infarct areas. (I) High magnification (63X) images showing PHH3 positive cells (Alexa 647, pseudo-color Magenta) and nuclei stained with DAPI (blue) highlighting their overlap and (II) corresponding overlay of GFP channel showing PHH3+/mGFP+ CMs.

To determine whether transfer of RBC-EVs miRNA cargo is associated with changes in the transcriptome of recipient cells, we studied the effect of RBC-EVs on a stable mT/mG HEK293 cell line we generated (see methods for details). We treated the mTmG reporter HEK293 cell line with EVs purified from the plasma of either EpoR-Cre or mTmG mice by differential gradient ultracentrifugation (**Figure S6A**, **B**). We observed Cre-mediated recombination (appearance of mGFP^+^ cells) only in the EpoR-Cre EV treatment group and not in mTmG EV-treated group (**Figure S1D, bottom panel**). Using flow sorting, we separated mTd^+^, mGFP^+^ and mTd^+^/mGFP^+^ sub-populations from mT/mG HEK293 treated with EpoR-Cre and mTmG EVs (**Figure 4C**). Overall, recombined cells (mGFP^+^ and mTd^+^/mGFP^+^) constituted 4.54±1.51% of the total population of flow sorted mT/mG HEK293 which were treated with EpoR-Cre EVs (**Figure S6C**).Targeted whole transcriptomic sequencing (Ion gene Studio S5 Next Generation Sequencing Systems) identified 143 genes that were expressed with at least 25 reads per million at a two-fold change between the two groups (mTd^+^ as one group vs mGFP^+^ and double positive mTd^+^/mGFP^+^ grouped together). The regulation of different molecular pathways by these 143 genes are shown by Ingenuity Pathway Analysis (IPA) (**Figure 4D**). Nine differentially expressed genes (COX8A, PPP2R5B, NDUFS7, CCNH, NDUFS6, SIRT2, ATP5L2, HDAC10 and E2F6) from relevant biological pathways (HDAC10 and SIRT2 in Adipogenesis pathway, E2F6 and PPP2R5B in Cell Cycle regulation by BTG family proteins and Role of CHK proteins in cell cycle checkpoint control, ATP5L2, COX8A, NDUFS6 and NDUFS7 in Oxidative Phosphorylation, CCNH, E2F6, PPP2R5B and HDAC10 in Cyclin and cell cycle regulation) were found to be changed in the same direction by QRT-PCR (**Table S2**). Taken together, our results suggest that the miRNA cargo of RBC-EVs can mediate transcriptional changes that may affect important cellular processes in recipient cells.

### Functional effects of RBC-EVs in ischemic heart failure

Cardiomyocytes have been shown to proliferate at low frequencies (Bergmann et al., 2009), and can re-enter the cell cycle after myocardial infarction (Li et al., 2013, Wang et al., 2017). Since the change in the transcriptome of HEK293 cells receiving RBC-EVs suggested a potential role of RBC-EV mediated signaling in regulation of cell cycle and cell division pathways, we explored the possibility of a similar role in the heart during post-MI cardiac remodeling. I/R was performed on EdU injected EpoR-Cre/mTmG mice and sacrificed after 4 weeks as a model of ischemic heart failure (**Figure S7A)**. Edu was incorporated in proliferating cells in both the spleen and thymus (**Figure S7B**), while in the heart, EdU^+^ cells were detected in higher numbers near the infarct zone compared to the remote zone in I/R mice (**Figure S7C**). While the overall frequency of EdU+ cardiomyocytes was low, consistent with other studies, we found that EdU^+^ cardiomyocytes were represented at a significantly higher (2-fold) proportion in GFP^+^ recombined cardiomyocytes compared to mTd^+^ cardiomyocytes (**Figure 4E**). Complementary to these results, immunostaining of heart sections from I/R EpoR-Cre/mTmG mice with another proliferation marker, Phospho-histone H3 (PHH3), showed a similar pattern of PHH3-immunoreactivity in mGFP^+^ cardiomyocytes (**Figure 4F,G**). Together our data suggest that RBC-EV mediated signaling by transfer of their miRNA content may alter transcriptional profiles of target cells and enhance the possibility of proliferation. Even though overall proliferative capacity in cardiomyocytes following MI is low, RBC-EV mediated signaling may play a role in priming cardiomyocytes for DNA synthesis and proliferation.

## Discussion

Extracellular vesicles (EVs) have recently been shown to constitute a novel mode of intercellular signaling by transferring their cargo molecules to recipient cells. RBCs are the most abundant cell type in the circulation, and RBC-EVs comprise a significant proportion of circulating EVs in human plasma (Danesh et al., 2014, Sender et al., 2016). RBC-EVs have recently been recognized as key regulators of various physiological and pathological processes, including coagulation (Danesh et al., 2014, Jank and Salzer, 2011, Kozuma et al., 2011), inflammation, atherosclerosis and thrombosis (Straat et al., 2016a, Buttari et al., 2015, Chung et al., 2007). In patients with heart failure, myocardial infarction or infections such as malaria, the levels of RBC-EVs are often found to be increased (Giannopoulos et al., 2014). Importantly, the complement activation at the site of inflammation makes RBC-EVs (Kuo et al., 2017, Karasu et al., 2018) poised to fuse with, and deliver their cargo to recipient cells. However, despite increased appreciation for their biological importance, mechanistic insight into the functional consequence of the RBC-EV: target cell fusion events are largely unknown.

To determine the *in vivo* targets and roles of RBC-EVs, we adapted a murine model previously described by the investigators to characterize the targets of hematopoietic cell-derived EVs, to specifically study the role of RBC-EVs in ischemic heart failure. The model, initially described by Ridder and colleagues (Ridder et al., 2014), utilizes a double-fluorescent Cre reporter that expresses membrane targeted tandem dimer tomato (mT) globally, but expresses membrane-targeted EGFP (mGFP) following Cre-mediated recombination. In the original study, the investigators found that when crossed with the vav-icre transgenic mice (Cre expression in hematopoietic cells), mGFP expression was noted in hematopoietic and immune cells. Interestingly Cre recombinase mRNA was packaged in EVs, and the functional Cre mRNA could be transferred to EV recipient cells, marking them as mGFP^+^ following recombination. We adapted this system to study RBC-EVs using EpoR-Cre mice. We found that RBCs, and their progenitors in the bone marrow, expectedly express mGFP, and EVs derived from RBC membranes are also mGFP positive. Platelets, that arise from the common megakaryocyte-erythrocyte progenitor (MEP) cells that express EpoR also show evidence of mGFP positivity; however, given that megakaryocytes do not express EpoR, while erythrocyte progenitors continue to express EpoR, the efficiency of recombination is likely much higher in RBC lineage as seen in our flow cytometry and microscopy data. Importantly, we found that Cre protein (and not mRNA) was packaged into RBC-EVs and was functional upon transfer to recipient cells not only in cell culture models but also *in vivo*. The fact that RBCs differ from other cells in not having a nucleus may account for the fact that Cre protein is packaged from RBC cytoplasm into the EVs. Identification of the RBC-EV target cells by the Cre-mediated fluorescence conversion of the target cells allowed us to study intercellular communication between RBCs and different cell-types both at baseline and in a validated ischemic HF model.

To address the issue of ectopic Cre expression or cell fusion accounting for recombination in cell types other than erythroid cells, we administered complement-generated RBC-EVs from EpoR-Cre mice into the mTmG mice and found the same pattern of recombined cells in these transfused mice as in the double transgenic EpoR-Cre/ mTmG mice, suggesting that the source of Cre in the recombined cells were in fact RBC-EVs. Multiple different cell types were targeted by RBC-EVs, and included cardiomyocytes, tubular epithelial cells in the kidney and splenic cells, but interestingly, not cells in the liver. Whether surface markers on RBC-EVs, or specific EV ‘receptors’ on these cells are responsible for the observed pattern is not clear at this point, as an understanding of the mechanisms of EV uptake has not yet been elucidated. Additionally, baseline levels of recombination are low, suggesting a modest to low level of signaling mediated by RBC-EVs in the absence of stress. Whether the uptake at baseline is a stochastic process or reflects a distinct functional hierarchy in same-type target cells remains to be determined.

Previous studies had suggested that EV-mediated signaling is implicated in disease pathogenesis, such as cancer cell metastasis, immune escape of tumor cells, and obesity-associated insulin resistance. Notably the RNA cargo of EVs have garnered significant attention, as active signaling moieties. In this regard, by account of the fact that the landscape of RBC-EVs is dominated by small RNAs, the RBC-EVs reflect this simpler composition compared to EVs from nucleated cells. Deep RNA sequencing from RBC-EVs from human cohorts revealed that the content of RBC-EVs is dominated by a small number of miRNAs, and that this content does not significantly alter in patients with type I myocardial infarction or heart failure. Using a computational approach, we found that the pathways predicted to be regulated by these miRNAs are implicated in cell cycle regulation, proliferation and oxidative metabolism. Using cellular RNA seq to distinguish cells targeted by RBC-EVs from non-targeted cells (in the HEK 293 mTmG reporters), we confirmed that differentially expressed genes belonged to pathways relevant to cell cycle, proliferation, and oxidative phosphorylation, some of which were individually validated by QRT-PCR. Future experiments using single cell nuclear RNAseq to delineate the transcriptome of targeted cells *in vivo* may reveal the effects of EV-mediated signaling in more granular detail but were beyond the scope of these initial studies.

Ridder et al had previously found that the stress of peripheral inflammation markedly increased the EV-mediated signaling between peripheral hematopoietic cells and neuronal cells. As RBC-EVs can be generated in large numbers at the site of complement activation, we hypothesized that intercellular signaling between RBCs and cardiomyocytes may increase in the setting of post-MI remodeling. Indeed, we found a significant increase in the number of recombined CMs in the IR model suggesting that similar to the model previously described from the vav-icre system, RBC-EV-mediated signaling can increase markedly following the stress of MI. The timing of these signaling events were not elucidated as we only examined the 4-week time point, at which time we found recombined cardiomyocytes not only at the peri-infarct zone (as expected), but also in remote myocardium. Similar observations have been made when using Cre-recombinase based tracing of EVs in the tumor microenvironment (Mateescu et al., 2017, Ridder et al., 2015), indicating the importance of the state of the uptaking cell, rather than the solely the presence of EVs. Whether the initial increase in RBC-EV uptake happens in the peri-infarct zone, where there is likely both compromise in the endothelial barrier, as well as complement activation, and then spreads to more remote zones will need more detailed time course experiments. Finally, we found that recombined cardiomyocytes were more likely to undergo DNA synthesis than non-recombined cardiomyocytes. We hypothesize that uptake of RBC-EVs may allow for transfer of miRNAs that may prime the cell for proliferation (associated with changes in expression of genes implicated in these processes in our cell culture experiments). However, we cannot prove that RBC-EV uptake does lead to the low-level increase in cardiomyocyte proliferation that has been noted by multiple groups in murine MI models. Additionally, in the absence of reagents that can inhibit or enhance uptake or release of EVs *in vivo* without affecting critical cellular processes, we cannot postulate on the importance of this signaling on disease pathogenesis. However, the changes in transcriptome and the phenotyping associations with markers of proliferation certainly makes a plausible case for an important role for EV-mediated signaling in post-MI remodeling.

Finally, the finding that microglia are targeted by RBC-EVs only following MI is intriguing. Microglia are the resident macrophages of the brain and are the first to respond to infection and injury (Saijo and Glass, 2011) and microglial activation has been reported in the brains of humans with heart disease (Streit and Sparks, 1997). Under inflammatory conditions, we have demonstrated that EVs released from hematopoietic cells can overcome BBB and transfer RNA to neural cells (Ridder et al., 2014). Studies by others have shown that EVs derived from RBCs, that are abundant in the toxic forms of α-synuclein protein, can cross the BBB under inflammatory conditions, and localize with brain microglia in patients with Parkinson’s disease (Matsumoto et al., 2017). Whether the increase in microglia targeted by RBC-EVs affect processes like depression or cognitive function that have increasingly been recognized as closely associated with ischemic heart disease is an intriguing hypothesis that merits future investigation.

Our study was limited by the inability to assess differences in the transcriptome profiling between recombined and non-recombined cardiomyocytes *in vivo*, as single-cell RNA-seq on cardiomyocytes remains challenging. Conventional methods are limited by the cell size, and large cells like cardiomyocytes have posed difficulties for single-cell RNA-seq, and conventional FACS-based methods for isolating cells from complex tissues, like the heart and the brain, often influence their mRNA composition. Recently developed techniques such as single nuclear RNAseq may be adapted in the future to answer this critical question. Secondly, it remains possible that only a proportion of RBC-EVs contain functional Cre, and that the cells marked by GFP positivity (and therefore Cre-mediated recombination) only constitute a proportion of cells that have taken up RBC-EVs; in this case we may be under-estimating the extent of intercellular communication mediated by this mechanism. In the case of liver, it is likely that RBC-EVs were internalized by the reticuloendothelial system (RE) system (Kupffer cells), and destroyed by the lysosomes, thus preventing the Cre to access the nucleus and convert the cells from tomato to GFP. Finally, while it is intriguing that only certain cell-types take up RBC-EVs, there remains a paucity of insight into the EV surface markers and cellular receptors that dictate this specificity.

In summary, our model is the first to demonstrate previously unreported communication between EVs released from RBCs and specific organs, both in health and inflammation. Furthermore, our data demonstrate functional changes in the transcript and phenotype of targeted cells, especially in the context of ischemic HF and post-MI remodeling. We hope that this model will be the basis of future investigations into the functional role of this novel signaling mechanism in altering cellular phenotype and modulating disease pathogenesis.

## Acknowledgements

SD was supported by grants from NIH (NCATS, UH3 TR000901, RO1 HL122547); SM DFG (MO2211-2) and Edinger Foundation.

## Author Contributions

A.D. and N.V. designed, conducted experiments and analyzed the data. A.M.S. performed flow cytometry experiments and analyzed data. I.K. and O.Z. conducted experiments. A.Y. analyzed RNA-seq data. F.G., S.L. and A.K. assisted in experiments. C.X. performed Sham and IR surgeries. R.S.R., S.S., V.T., G.L. and J.T. performed experiments. L.L. assisted with sequencing experiments. S.M., I.G. and S.D. conceived the project, and wrote the manuscript.

## STAR Methods

### Collection of CFP for isolation of EVs

Mice (8-10 weeks old) were anaesthetized using 2.4% isoflurane/97.6% oxygen and placed in a supine position on a heating pad (37° C). Cervical dislocation was then conducted in the mice and blood collected from intracardiac puncture using 27mm gauge needle. The blood was collected in Microvette® Micro tube, 1.3ml, screw cap, EDTA tubes and centrifuged at 1000g for 15 minutes. The supernatant was collected in fresh Eppendorf tubes, leaving the buffy coat and the RBC fraction. The buffy coat was discarded, and the RBC fraction was transferred in separate Eppendorf tubes for generation of complement-mediated RBC-EVs. The supernatant from first spin was centrifuged again at 1500xg for 10 minutes. Then supernatant was Cell Free Plasma (CFP), which was taken in fresh Eppendorf tubes leaving the pellet behind and processed for EV isolation.

All human patient-related investigations were conducted in conformation with the principles outlined in the Declaration of Helsinki and duly approved by the relevant institutional review committee (Massachusetts General Hospital). Written informed consent were obtained from all participants before their enrollment in the study. Human venous blood was drawn from healthy volunteers, patients with Type I MI, or with chronic heart failure with the help of 21G needle under aseptic condition and the blood was collected in EDTA-containing Vacutainer Tubes. The blood was centrifuged at 1000xg for 15 minutes. The supernatant was collected in fresh 15 ml plastic tubes, leaving the buffy coat and the RBC fraction. The supernatant from first spin was centrifuged again at 1500xg for 10 minutes and the supernatant was Cell Free Plasma (CFP), which was taken in fresh tubes leaving the pellet behind and processed for EV isolation.

### Complement mediated generation of RBC-EVs

Ten microlitres of RBC from tubes fraction of mice blood was resuspended in in HBSS++ buffer (1:2 by volume) and centrifuged in table top centrifuge to form a pellet. Supernatant HBSS++ is discarded, pellet is resuspended in HBSS++ and the process is repeated twice. C5b,6 solution (0.18μg/ml final concentration in HBSS++) was added, vortexed to mix properly and put on a slow shaker at room temperature for 15 minutes. C7 (reconstituted to a final concentration of 0.2μg/ml in HBSS++) was added and put on a slow shaker at room temperature for 5 minutes. C8 (to a final concentration of 0.2μg/ml in HBSS++) and C9 (to 0.45μg/ml in HBSS++) was simultaneously added to the solution and incubated at 37°C for 30 minutes. After centrifuging the solution at 2500×g for 10 minutes, the EV-containing supernatant was collected in a new tube.

### Isolation of EVs from CFP

Isolation of EVs from CFP (mice and human) was performed using a well-described protocol (Li et al., 2018) with modifications. 3 ml of CFP was resuspended in 5ml of filtered PBS and underlaid with 2 ml of 60% iodixanol and centrifuged at 100,000xg for 2 hours. The top 7 ml of supernatant was discarded and the 40% iodixanol with EVs was underlaid over a column gradient consisting of 20%, 10% and 5% iodixanol (suspended in sucrose) and centrifuged at 100,000xg for 18 hours. Among the 12 fractions, fraction 6-8 was primarily used for our experiments. For the functional experiments with EVs (treating HEK293 cells) an extra step of overnight dialysis of the EVs suspended in iodixanol was done with Spectra-Por® Float-A-Lyzer® and finally resuspended in filtered PBS.

### Nanoflow cytometry

The principles and detailed workflow of Nanoflow cytometry for characterization of EVs was done as per our previously published work (Bei et al., 2017). For CFP from EpoR-Cre, EpoR-Cre/mTmG and mTmG mice, the gated population for EVs was further analyzed for detection of dTomato and GFP. The quantification of EVs (fluorescent and non-fluorescent) was also done as per our previous published work (Bei et al., 2017).

### Primary dermal fibroblast and mTmG HEK293 cell lines

Isolation and culture of primary dermal fibroblast was done using a previously described protocol (Seluanov et al., 2010). On attainment of confluence, the cells were trypsinized and passaged in 6 well tissue culture dishes (1:5) and kept in tissue culture incubator for 24 hours. Subsequently, the dermal fibroblasts were treated with 150 microliters of plasma EVs (which were isolated through sequential density ultracentrifugation as mentioned above) in each well of 6 well dish. The cells were put in the incubator for 72 hours, after which live cell imaging with fluorescence microscopy was done on them.

Cryopreserved HEK 293T cells were seeded in a Fibroblast growth and culture media (FGCM) (DMEM, 5 %FBS, 1% PS, pH 7.4). On attainment of 70-80% confluency, the cells trypsinized and split (1:10) into 6-well cell culture plates and cultured in FCGM. pCA-mTmG and pTriEx-1plasmids was inoculated in separate LB Agar (with Ampicillin) plates and the colonies were amplified, overnight, in liquid LB broth with Ampicillin. The DNA from both the plasmids were extracted using Qiagen Plasmid Maxi Kit and stored at −80*C in TE buffer. The adjusted volume of DNA from pCA-mTmG plasmid were separately solubilized in Lipofectamine 3000 and treated in separate well of 6 well dishes (2500ng DNA per well, 5 microliters of P3000 reagent per well, 3.75 microliters of Lipofectamine 3000 reagent per well, dissolved in 250 microliters of Opti-MEM™ I Reduced Serum Medium per well). After incubation in FCGM media for 12 hours, the antibiotic in media was changed from PS to Blasticidin for antibiotic selection of HEK293 cells. The cells were periodically checked under fluorescence microscope to check for the fluorescence and confluency of mTmG HEK293 cells. The media was changed after every 24 hours to get rid of the dead and floating HEK293 cells and the cell line was passaged and the clone that showed the maximum integration of plasmid containing DNA was maintained for future use as a stable mTmG HEK293 cell line. The mTmG HEK293 cells were also treated with 100 microliters of plasma EVs from EpoR-Cre mice (which were isolated through sequential density ultracentrifugation as mentioned above) in each well and the wells of mTmG HEK293 cells which were treated with DNA from pTriEx-1plasmids (Lipofectamine 3000) was considered as positive control.

### FACS analysis and sorting

Adult mouse CMs were isolated from Sham or IR operated mice’s hearts using a Collagenase B (Roche, 11088807001), Collagenase D (Roche, 11088858001) and Protease XIV (Sigma-Aldrich, P5147) based digestion, following the isolation steps described by Ackers-Johnson et al. Circ Res. 2016; 119:909-920. For live-cell sorting and quantitative analysis, adult mouse CMs and HEK293 cells were stained with DAPI (BioLegend, 422801) for 30min in the dark, sorted using a BD FACSAria and analyzed by FlowJo.

### RNA Isolation, Pre-amplification and Quantitative Real Time PCR (qRT-PCR)

Total RNA was extracted from EV samples and FACS sorted HEK293 cells using mirVana Paris RNA and Native Protein Purification Kit with modifications suggested in an established protocol (Burgos and Van Keuren-Jensen, 2014) and further concentrated and purified using RNA Clean and Concentrator-5 with Dnase I Set. The RNA was then measured using 4200 TapeStation Instrument using High Sensitivity RNA Screen Tape. For mRNA qPCR cDNA was created from equivalent RNA from each sample with High Capacity cDNA reverse Transcription Kit in Biorad CFX384 qPCR System. For detection of Cre-recombinase mRNA, amplification and detection of specific products was done using the ExiLENT SYBR® Green master mix in QuantStudio 6 Flex Real-Time PCR System till 45 cycles. Any CT equal or above 40 was considered unreliable and the mRNA was considered not be absent in that sample. For the FACS sorted HEK293 cells, after the creation of cDNA, the cDNA was then pre-amplified using Taqman PreAmp Master Mix for 20 cycles using the manufacturer’s protocol. Further amplification and detection of specific products were performed with Taqman primers and TaqMan Gene Expression Master Mix in QuantStudio 6 Flex Real-Time PCR System till 45 cycles. Any CT equal or above 40 was considered unreliable and the mRNA was considered not be absent in that sample. For HEK293 cells, GAPDH was considered as internal control for the entire experiment.

For microRNA, cDNA from total RNA, after clean-up and concentration, was constructed using Universal cDNA Synthesis Kit II in Biorad CFX384 qPCR System (maximum possible RNA input in 20 microliters reaction). Amplification and detection of specific products was done using the ExiLENT SYBR® Green master mix along with Exiqon miR LNA PCR primer set UniRT, specific to each miRNA, in QuantStudio 6 Flex Real-Time PCR System till 45 cycles.

### RNASeq of Human Plasma Samples

RNA was isolated from 1 mL plasma using the miRCURY RNA Isolation kit for Biofluids (Exiqon) with modified protocol. Libraries were constructed and amplified from approximately 3 ng RNA using the NEXTflex small RNA-Seq Kit V3 for Illumina Platforms (Bioo Scientific, a PerkinElmer company). Size selection of libraries was performed by gel electrophoresis on a 10% TBE gel (Invitrogen) with excision of the 140 to 160 nucleotide (nt) bands (corresponding to 21–40ntRNA fragments). Libraries were sequenced on an Illumina HiSeq platform for single read 50 cycles at the NextGen Sequencing Core at Massachusetts General Hospital.

### RNA-Seq data analysis

The raw sequence image files from the Illumina HiSeq 2500 or Illumina MiSeq in the form of bcl are converted to the fastq format using bcltofastq v1.8.4 and checked for quality to ensure the quality scores do not deteriorate drastically at the read ends. The adapters from the 3’ end were clipped using cutadapt v.1.10 (Martin, 2011) (http://cutadapt.readthedocs.io/en/stable/guide.html). Reads shorter than 15nts are discarded and after adapter trimming, the 3’ bases below a quality score of 30 are trimmed as well. Reads that arise from human rRNA and contamination from library preparation protocols are removed before they are mapped to the human genome. The reads are first mapped to a library of UniVec contaminants, a collection of common vector, adapter, linker and PCR primer sequences collated by the NCBI. They are then mapped to human rRNA sequences obtained from NCBI The reads are mapped to the rRNA and UniVec sequences using Bowtie2 (Langmead et al., 2009) and those that map are removed from the analysis. The alignment to the human genome and transcriptome takes place in two stages. First, the rRNA and UniVec free reads are mapped to the human genome (hg19) using STAR (Dobin et al., 2013). The reads that map to the genome are then mapped to the human transcriptome. Also, the reads that are not mapped to the human genome are mapped to the human transcriptome, The library for the human transcriptome is built by concatenating miRNAs and hairpins from miRBase 21 (Kozomara and Griffiths-Jones, 2011), tRNAs from gtRNAdb (Chan and Lowe, 2009), piRNAs from piRBase v1.0 (Zhang et al., 2014), protein-coding, non-coding and other RNA sequences from ENSEMBL 75. The STAR alignment is performed end to end with a single mismatch allowed while mapping and each read is allowed to multimap to at most 40 RNA annotations. Here, there is no mismatch allowed and each read is allowed to multimap to at most 40 RNA annotations. The miRNA expression is calculated as the number of miRNA reads mapped per million reads mapped to the human transcriptome, referred to as RPM in short. In these experiments, input RNA reads from each group (control, MI and heart failure) were analyzed as pool samples due to low overall reads from the isolated EV populations.

### Pathway analysis

The targets of these top 20 miRNAs were identified using TargetScan v 7.1 and all target mRNAs with a cumulative context score less than 0.2 were included for Gene Ontology analysis using DAVID (Huang da et al., 2009). The p-values represented in Figure 4B are based on a Fisher’s Exact test.

Genes that were expressed with at least 25 RPM at a two-fold change between the two groups (mTd^+^ as one group vs mGFP^+^ and mTd^+^/mGFP^+^ grouped together) were introduced into Ingenuity Pathway Analysis (IPA). The p-values represented in Figure 4D are based on a Fisher’s Exact test.

### Animal care and use

All animal studies were approved by the Massachusetts General Hospital Animal Care and Use Committee and under the guidelines on the use and care of laboratory animals for biomedical research published by National Institutes of Health (No. 85-23, revised 1996). The generation of EpoRCre mice (Heinrich et al., 2004) and ROSAmT/mG (mT/mG) (Muzumdar et al., 2007) have been previously described. Wild-type (WT) C57BL6 mice were received from Jackson Laboratory. Maintenance of 12/12 hour light-dark cycle was done for all mice. Mice were fed on normal chow diet. Male/female mice of 8-12 weeks were fed ad libitum.

For the transfusion experiments, mice were injected intraperitoneally with a single dose of 700 µl of complement generated RBC-EVs (RBC-EVs generated with complement suspended in the complement soup). The mice were caged in barrier facility and monitored for 7 days before they were sacrificed, and blood was collected through intracardiac organs (heart, lungs, spleen, kidney, liver and brain) were harvested.

For I/R surgery, the left anterior descending artery (LAD) was ligated with 7-0 silk. After five minutes of ischemia, 50 ml of fluorescent microspheres (10 mm FluoSpheres, Molecular Probes) were injected into the LV cavity. Following 30 min LAD occlusion, the LAD ligature was released, and reperfusion was confirmed visually. After 4 weeks of reperfusion, mice were sacrificed, and organs were harvested for analyses. For the sham surgery, the animals were also intubated, and underwent sternal skin incision without a thoracotomy to minimize the effect of inflammation noted after thoracotomy (Mitsos et al., 2009).

Echocardiography was conducted on conscious mice using a GE Vivid7 with i13L probe (14 MHZ) as described previously (Das et al., 2012). Apart from the 2D guided M-mode images of short axis at the papillary muscle, parasternal long-axis views and short-axis views were also recorded. The average of at least ten measurements was used for every data point from each mouse.

Cardiomyocyte (CM) isolation from I/R and sham mice was done using a previously published protocol (Ackers-Johnson et al., 2016). Flow sorting of CM population was done and the enrichment of the viable CMs through this preparation was tested by qPCR of commonly expressed genes in CMs compared to the other fraction from the preparation. CM isolated from EpoR-Cre/mTmG mice after I/R surgery were further sorted based on their fluorescence.

EdU (50 mg/kg, subcutaneously) in EpoR-Cre/mTmG mice, after I/R, was injected every alternate day for the first 14 days after reperfusion and the animals were euthanized after 6 weeks. Sham-operated EpoR-Cre/mTmG mice (without thoracotomy) served as controls.

### Western Blot, Immunochemistry and Immunofluorescence Staining

Isolated EVs from EpoR-Cre and mTmG mice with ExoQuick Plasma prep and Exosome precipitation kit were resuspended in 30 µlof filtered PBS. Standard western blot technique and analysis was done on equal amount of protein using Anti-Cre Recombinase (Abcam, Cat#: ab24607, Dilution 1:200), CD9 (Biolegend, Cat#: 124802, Dilution 1:200) and Ter 119 (Thermo Fisher Scientific, Cat#: 14592181, Dilution 1:200) antibodies. The EVs from human plasma isolated through C-DGUC (individual fraction of 1 ml) was again ultracentrifuged for 2 hours for precipitation of EVs and subsequently resuspended in 30 µlof filtered PBS before Standard western blot technique and analysis was done on equal amount of protein using CD9 (Biolegend, Cat#: 312102, Dilution 1:1000) and Anti-Alix (Biolegend, Cat#: 634501, Dilution 1:1000) antibodies. The exposure time to detect individual antibodies is included in the figure legend.

Organs (heart, lung, liver, spleen, kidney and brain) were harvested from individual mice and fixed in Paraformaldehyde (4%), overnight. The organs were then cryoprotected with 20% sucrose solution till the organs start sinking in the solution. Then they were thoroughly washed with PBS and mounted in O.C.T before cryo-sectioning into 10 micron tissue sections. The slides were mounted in DAPI medium before confocal microscopy. For the brain, sagittal sections were cut on a vibrotome and kept in PBS at 4°C.

For PHH3 and Iba staining, heart and brain sections, respectively, were washed with PBS and acetone/methanol solution before blocking with 3% (w/v) BSA in PBS and TritonX/BSA/PBS and then incubated with primary antibodies applied (PHH3 Cell Signaling Technology, Catalog Number: 3377S, diluted 1:250 and Iba1, Fujifilm Wako Pure Chemical Corporation, Catalog Number 01919741, diluted 1:1000) and for Anti-NeuN antibody (Fujifilm Wako Pure Chemical Corporation, Cat#:01919741) overnight at 4° C. The sections were sequentially washed in PBS/Triton X and treated with secondary antibody (1:1000) for 1 hour, before mounting the sections with DAPI for confocal microscopy.

For EdU staining, after the sections were treated as mentioned above, before the incubation with primary antibody. EdU detection was conducted using manufacturer’s protocol and DNA staining was done using Hoechst 33342 using the same protocol. For

### Isolation of CD235a positive EVs from patient plasma

CD235a antibody (1µg/µl, Thermo Fisher Scientific, Cat#:MA512484) was added along with 5 mg of dynabeads and incubated overnight at 4° C. After washing the beads with Native Lysis Buffer twice and resuspension in 500 µl of Native Lysis Buffer along with RNAseOUT, DTT and PIC solution, and 30µl of CD235a coated beads was added to individual patient plasma sample and rotated at 4° C for 2 hours. After magnet induced retrieval of CD235a bound EVs in the dynabeads, the supernatant is discarded, and reverse crosslinking was done following manufacturer’s protocol for retrieval of CD235a^+^ EVs.

## Supplemental Information

### Supplemental titles and legends

**Figure S1. Baseline characterization of RBC-EVs in transgenic murine model**

(A) Flow cytometry scatter plots of isolated RBCs from peripheral blood of EpoR-Cre/mTmG mice (after removal of buffy coat) shows the presence of GFP+/Ter 119+ population (Q4, background (no fluorescence), Q1, Ter 119 (Brilliant Violet 421), Q3 as mGFP+(green) and Q2 as Ter 119+/mGFP+(double-positive)). Ter 119- but GFP+ cells represent contaminating platelets in the RBC preparation.

(B) Nanoflow cytometric characterization of CFP of EpoR-Cre/mTmG mice, showing the absence of any fluorescence in this group of mice. The auto fluorescence from the Suspension Buffer (Phosphate Buffer Saline) has been accounted for, as background. Shown are Td on the y-axis and GFP in the x-axis and divided in four quadrants (B--, background (no fluorescence), B-+ mTd+(red), B+- as mGFP+(green) and B++ as mTd+/mGFP+(double-positive)).

(C) Quantification of EpoR-Cre/mTmG CFP EVs (expressed as percentage of total fluorescent EVs): mTd+ (25.47±10.84%), mGFP+ (44.47±6.71%) and mTd+/mGFP+ (24.34±4.86) population (n=7). Data is presented as Median±SEM.

(D) High magnification fluorescent microscopy (40X) of fixed and mounted primary dermal fibroblasts from mTmG mice (Top) and low magnification confocal microscopy (20X) of fixed and mounted reporter mTmG HEK293 cells (Bottom) treated with (i) Adeno-Cre vector (positive control) (ii) mTmG RBC-EVs (iii) EpoR-Cre RBC-EVs respectively, to observe the pattern of change in membrane-bound fluorescence (n=3 biological replicates in each group).

**Figure S2. In-vivo transfer of functional Cre-recombinase through RBC-EVs**

(A) Schematic representation of experimental design for intraperitoneal injection in WT mice with complement-generated EpoR-Cre/mTmG RBC-EVs. Presence of mGFP+ EVs in CFP in WT mice (n=3) provides proof-of-concept about the route of administration of RBC EVs.

(B) Low magnification fluorescence microscopy (20X) of paraffin embedded sections of liver and brain from mTmG, EpoR-Cre/mTmG and mTmG mice transfused with EpoR-Cre RBC-EVs to characterize organ-specific tissue architecture. Corresponding, high magnification (63X) confocal microscopy of fixed frozen section of liver and brain from mTmG, EpoR-Cre/mTmG and mTmG mice transfused with EpoR-Cre RBC-EVs to map their fluorescence profile (on mTd and mGFP) at baseline and after transfusion. The findings were separately verified using biological replicates (n=4 for mTmG, n=6 for EpoR-Cre/mTmG and n=4 for mTmG mice transfused with EpoR-Cre RBC-EVs).

(C) Box and whisker plot demonstrating the count profile of EVs from CFP of mTmG mice transfused with EpoR-Cre, according to their fluorescence from Nanoflow cytometry. mTd+ EVs, mGFP+ EVs and mTd+/mGFP+ EVs constitute 51.15±15.07%, 15.99±15.78% and 6.54±13.14% of the total fluorescent EVs (n=4). Data is presented as Median±SEM.

**Figure S3. Count profile of recombination in organs receiving Cre-containing EVs by confocal microscopy**

Box and whisker plot showing the recombination profile (mGFP+ cells expressed as a percentage of total cells) of Baseline EpoR-Cre/mTmG mice and mTmG mice transfused with EpoR-Cre RBC EVs: kidney (9.76±1.49% vs 8.97±0.82%), heart (7.14±3.61% vs 10.31±1.45%), lung (10.44±2.46% vs 4.27±0.56%) and spleen (28.84±3.37% vs 11.67±2.11) (n=3 mice for Baseline and n=4 mice for transfused group, 40 high power filed per organ per mice). Data is presented as Median±SEM.

**Figure S4. Profiling of RBC-EVs post I/R**

Nanoflow cytometric characterization of CFP of EpoR-Cre/mTmG mice in Sham and I/R groups. The auto fluorescence from the Suspension Buffer (Phosphate Buffer Saline) has been accounted for, as background. Shown in the Sham group are Td on the y-axis and GFP in the x-axis and divided in four quadrants (B--, background (no fluorescence), B-+ mTd(red), B+- as mGFP+(green) and B++ as mTd+/mGFP+(double-positive)) and in I/R group are Td on the y-axis and GFP in the x-axis and divided in four quadrants (C--, background (no fluorescence), C-+ mTd+(red), C+- as mGFP+(green) and C++ as mTd+/mGFP+(double-positive)).

(A) Box and whisker plot quantifying EVs from CFP of EpoR-Cre/mTmG mice after Sham and I/R (67.15±25.59 vs 116.63±47.62, p=0.37, t-test, two tailed) (n=7 for Sham and n=6 for I/R). Individual CFP was run on Nanoflow cytometer and geometric means for X-axis and Y-axis for all EVs were obtained for each CFP of each group. The arithmetic mean of the geometric means of X-axis and Y-axis provided the count of all EVs from CFP of individual EpoR-Cre/mTmG mouse from each group.

(B) Box and whisker plot quantifying recombination profile of EVs from CFP of EpoR-Cre/mTmG mice after Sham and I/R (84.69±3.74% vs 78.61±3.19, p=0.12, t-test, two tailed) (n=7 for Sham and n=6 for I/R). Individual CFP was run on Nanoflow cytometer and geometric means for X-axis and Y-axis for individual group of EVs (based on fluorescence) were obtained for each CFP of each group. The arithmetic mean of the geometric means of X-axis and Y-axis provided the total count of EVs (of each individual fluorescence) from CFP of individual EpoR-Cre/mTmG from each group. The recombination profile is measured as the fraction of count of mGFP+ and mTd+/mGFP+ (combined) out of total EVs for an individual EpoR-Cre/mTmG mouse and expressed as percentage.

(C) Box and whisker plot showing the confocal microscopy count profile of recombination (mGFP+ cells expressed as a percentage of total cells) of Sham EpoR-Cre/mTmG mice and I/R EpoR-Cre/mTmG mTmG mice and EpoR-Cre/mTmG mice at baseline: kidney (5.97±0.56% vs 5.94±1.23% vs 9.76±1.49%), lung (8.85±0.76% vs 9.76±1.87% vs 10.44±2.46%) and spleen (31.93±4.19% vs 26.56±4.06% vs 28.84±3.37%) (n=4 mice for Sham group and n=5 mice for I/R group and n=3 mice for Baseline group, 40 high power filed per organ per mice). Data is presented as Median ± SEM.

**Figure S5. Transfer of Cre-recombinase through RBC-EVs during post infarct cardiac remodeling**

(A) Schematic representation of mTmG mice with intravenous transfusion of EpoR-Cre RBC EVs during I/R.

(B) Representative flow cytometry scatter-plots of DAPI- (viable) CMs from Control and I/R mice comparing the mGFP+ CMs between both groups (n=3 in each group). Shown are Td on the y-axis and GFP in the x-axis and divided in four quadrants (Q4, background (no fluorescence), Q1 mTd+(red), Q3 as mGFP+(green) and Q2 as mTd+/mGFP+(double-positive)). mTmG mice with intraperitoneal transfusion of EpoR-Cre RBC EVs act as Control in these experiments.

(C) Low magnification fluorescence microscopy (20X) of isolated CMs in mTmG mice with intravenous transfusion of EpoR-Cre RBC EVs during I/R (n=3 in each group).

**Figure S6. *In-vitro* transfer of functional Cre-recombinase through RBC-EVs**

(A) Schematic representation of isolation of EVs into 12 fractions, from mouse CFP using C-DGUC

(B) Detection of EV markers Alix and CD9 in the 12 fractions from C-DGUC through Western Blot. Molecular weight markers are indicated. The exposure time to detect Alix and CD9 was 200 secs.

(C) Box and whisker plot quantifying the frequency of recombination (mGFP+ and mTd+/mGFP+ combined, expressed as percentage of total fluorescent living cells) of mTmG HEK293 cells treated with EpoR-Cre RBC EVs (4.54±1.51%, n=9, 6 well-cell culture dish representing one individual event) (n=9 biological replicates). Data is presented as Median±SEM.

**Figure S7. Communication of RBC-EVs with proliferating cardiomyocytes**

(A) Box and whisker plot quantifying Fractional Shortening in Sham and I/R group of EpoR-Cre/mTmG mice, injected with EdU (50.3±3.09% vs 24.9±2.5%, p=0.005; n=3 for each group). Data is presented as Median±SEM.

(B) Processing of fixed and frozen sections of EpoR-Cre/mTmG mice spleen and thymus with Click-iT EdU Alexa 647 Imaging kit (to highlight proliferating cells) and confocal microscopy of the same. (I) High magnification (63X) imaging characteristics of nuclei (DAPI) and EdU (pseudo-color Magenta) to highlight their overlap and (II) corresponding overlay of GFP channel to characterize EdU+/mGFP+ individual cells (n=4 for I/R, n=3 for Sham).

(C) Processing of fixed and frozen sections of EpoR-Cre/mTmG mice heart with Click-iT EdU Alexa 647 Imaging kit (to highlight proliferating cells) and confocal microscopy of Infarct and Peri-infarct zone of I/R mice. Low magnification (20X) imaging characteristics of nuclei (DAPI) and EdU (pseudo-color Magenta) and corresponding overlay of GFP channel to characterize EdU+/mGFP+ CMs (n=4 for I/R, n=3 for Sham).

### Supplemental tables

**Table S1:**
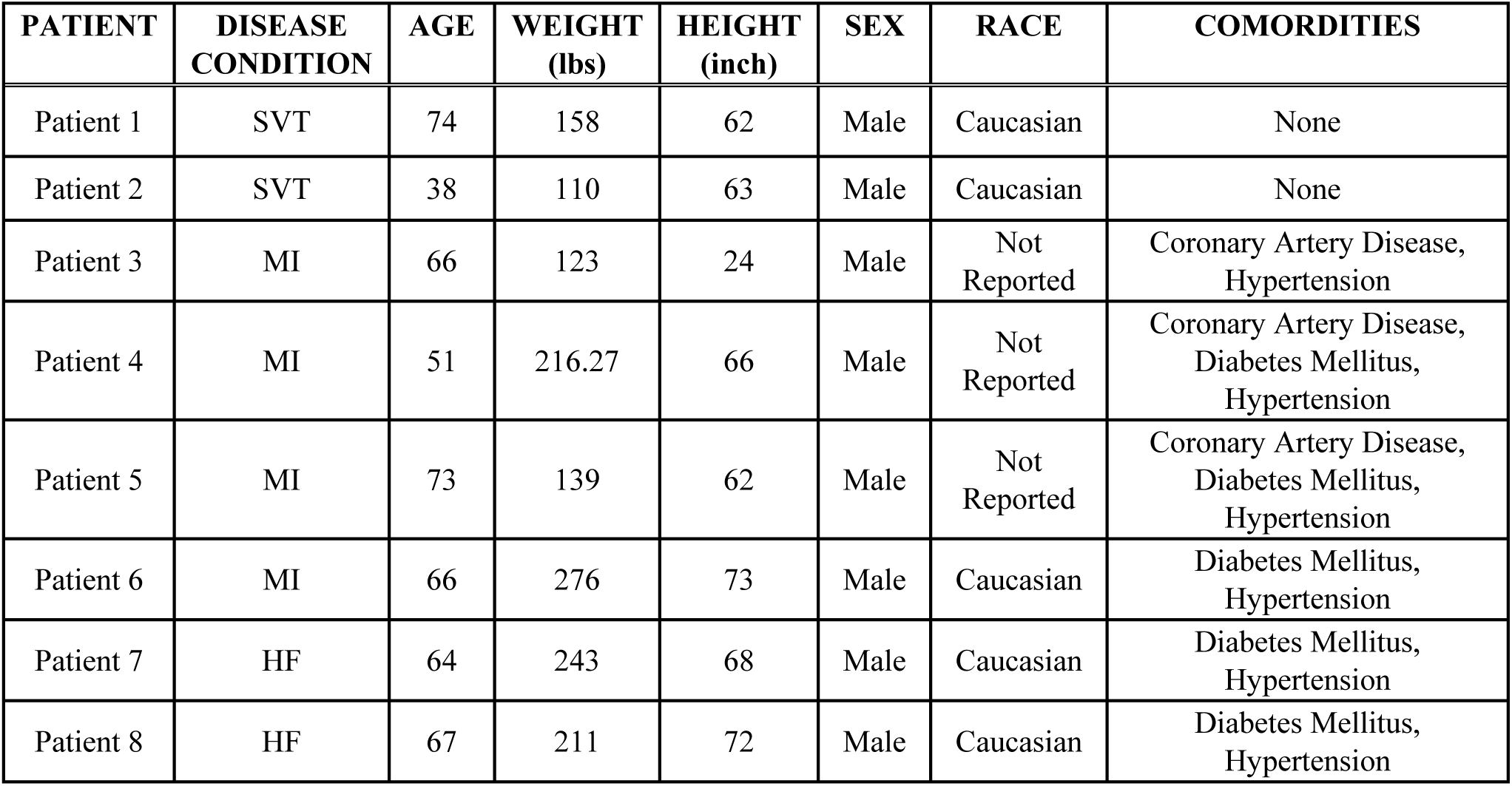
Clinical characterization of patients, whose plasma was used to characterize RBC-EVs RNA content. SVT: Supraventricular Tachycardia; MI: Myocardial Infarction; HF: Heart Failure

| **PATIENT** | **DISEASE**<br>**CONDITION** | **AGE** | **WEIGHT**<br>**(lbs)** | **HEIGHT**<br>**(inch)** | **SEX** | **RACE** | **COMORDITIES** |
| --- | --- | --- | --- | --- | --- | --- | --- |
| Patient 1 | SVT | 74 | 158 | 62 | Male | Caucasian | None |
| Patient 2 | SVT | 38 | 110 | 63 | Male | Caucasian | None |
| Patient 3 | MI | 66 | 123 | 24 | Male | Not<br>Reported | Coronary Artery Disease,<br>Hypertension |
| Patient 4 | MI | 51 | 216.27 | 66 | Male | Not<br>Reported | Coronary Artery Disease,<br>Diabetes Mellitus,<br>Hypertension |
| Patient 5 | MI | 73 | 139 | 62 | Male | Not<br>Reported | Coronary Artery Disease,<br>Diabetes Mellitus,<br>Hypertension |
| Patient 6 | MI | 66 | 276 | 73 | Male | Caucasian | Diabetes Mellitus,<br>Hypertension |
| Patient 7 | HF | 64 | 243 | 68 | Male | Caucasian | Diabetes Mellitus,<br>Hypertension |
| Patient 8 | HF | 67 | 211 | 72 | Male | Caucasian | Diabetes Mellitus,<br>Hypertension |

**Table S2:** Tabular representation of the fold change of nine differentially expressed genes selected from whole transcriptomic sequence of top relevant biological pathways and their validation. Fold change of nine differentially expressed genes from whole transcriptomic sequence analysis of recombined HEK293 cells (mGFP+ /mTd+ HEK293 cells). These genes belong to top relevant biological pathways: HDAC10 and SIRT2 in Adipogenesis pathway, E2F6 and PPP2R5B in Cell Cycle regulation by BTG family proteins and Role of CHK proteins in cell cycle checkpoint control, ATP5L2, COX8A, NDUFS6 and NDUFS7 in Oxidative Phosphorylation, CCNH, E2F6, PPP2R5B and HDAC10 in Cyclin and cell cycle regulation. Validation using qPCR and mutual concordance of the genes with two separate techniques.

| SEQUENCING DATA |  |  | Real-time PCR |
| --- | --- | --- | --- |
| GENE NAME | NCBI NAME | GREEN/RED LOG FOLD CHANGE | CONCORDANCE |
| ATP5L2 | ATP_synthase__H_transporting__mitochondrial_Fo_complex__subunit_G2 | 1.624131414 | Yes |
| E2F6 | E2F_transcription_factor_6 | 1.452054235 | Yes |
| HDAC10 | histone_deacetylase_10 | 1.19828344 | Yes |
| PPP2R5B | protein_phosphatase_2__regulatory_subunit_Bp__beta | -1.381467967 | Yes |
| CCNH | cyclin_H | -1.540152179 | Yes |
| COX8A | cytochrome_c_oxidase_subunit_VIIIA__ubiquitous | -1.495647613 | Yes |
| NDUFS7 | NADH_dehydrogenase__ubiquinone__Fe-S_protein_7__NADH-coenzyme_Q_reductase | -1.563667722 | Yes |
| NDUFS6 | NADH_dehydrogenase__ubiquinone__Fe-S_protein_6__NADH-coenzyme_Q_reductase | -1.629479318 | Yes |
| SIRT2 | sirtuin_2 | -1.707010076 | Yes |

