## Supplementary Figures for "Red blood cell-derived extracellular vesicles mediate intercellular communication in ischemic heart failure"

Supplemental Fig. 1

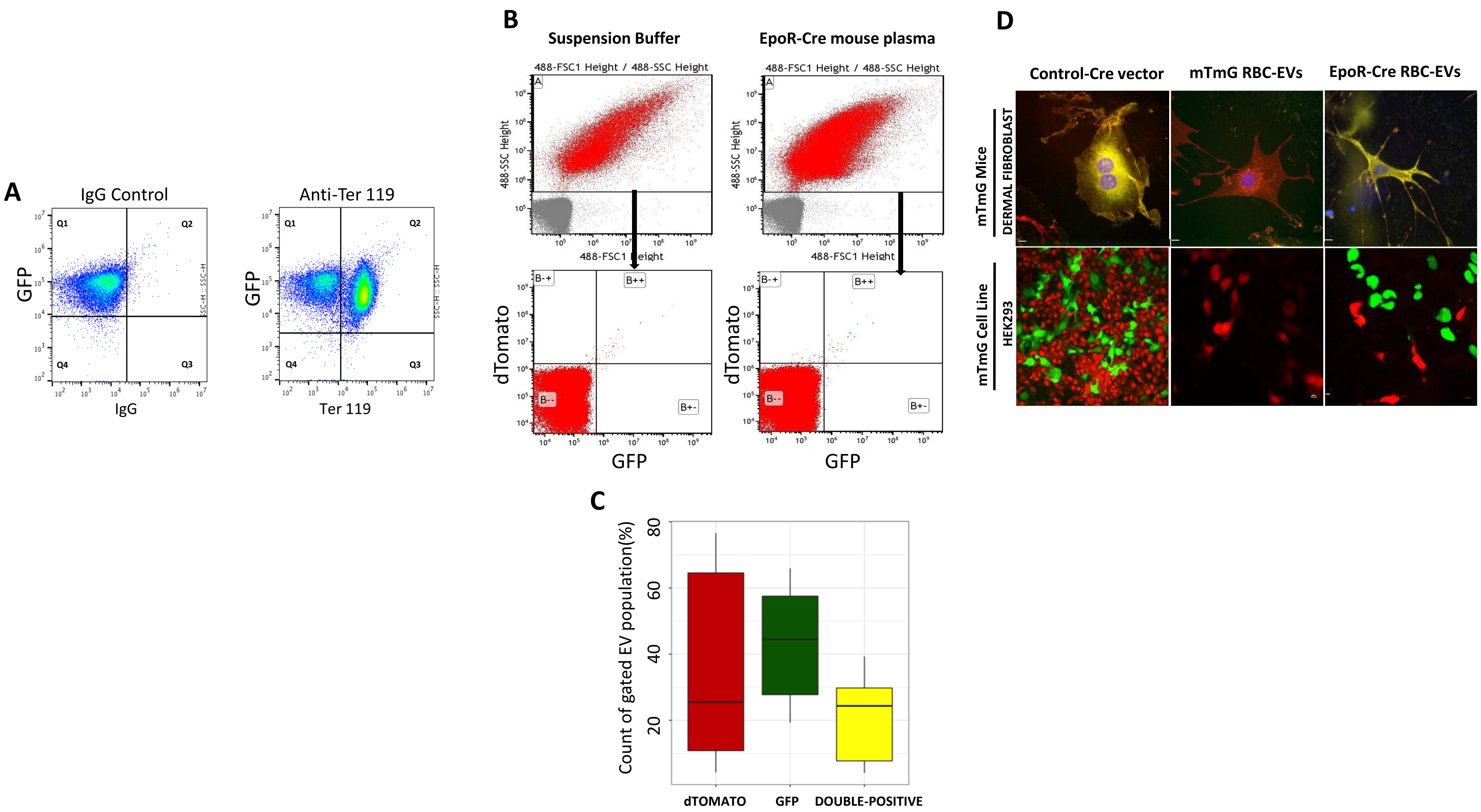

Supplemental Fig. 2

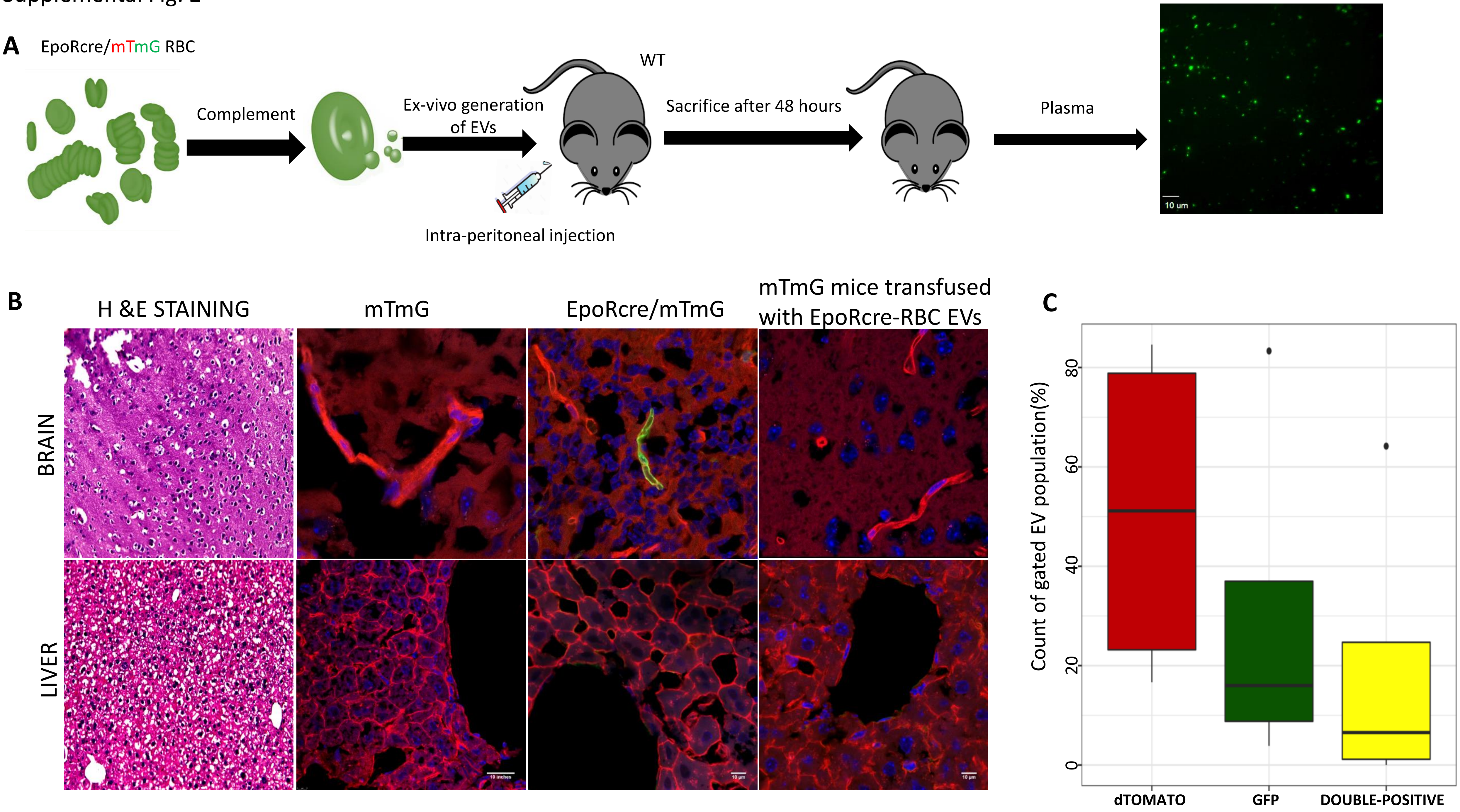

Supplemental Fig. 3

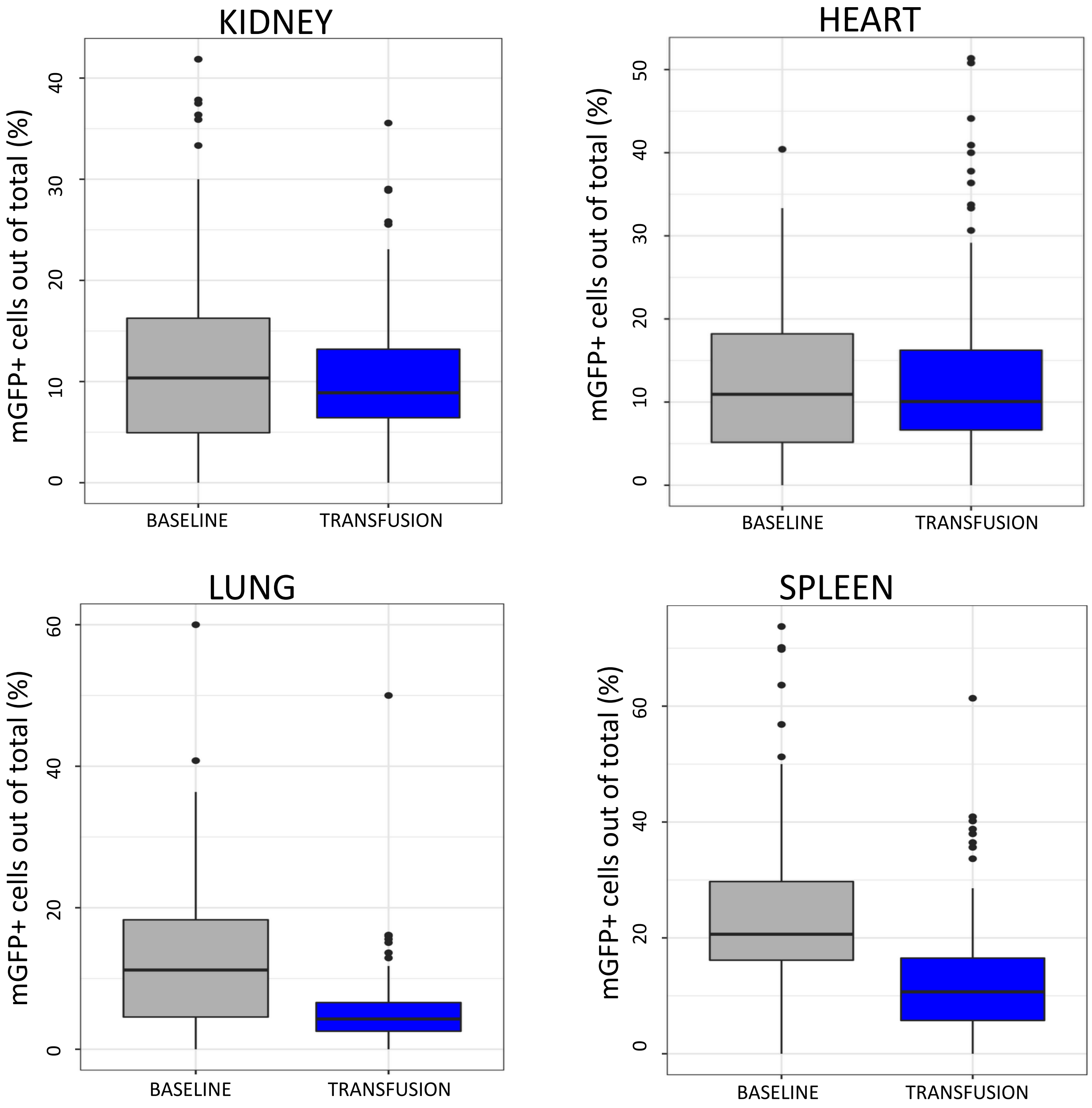

Supplemental Fig. 4

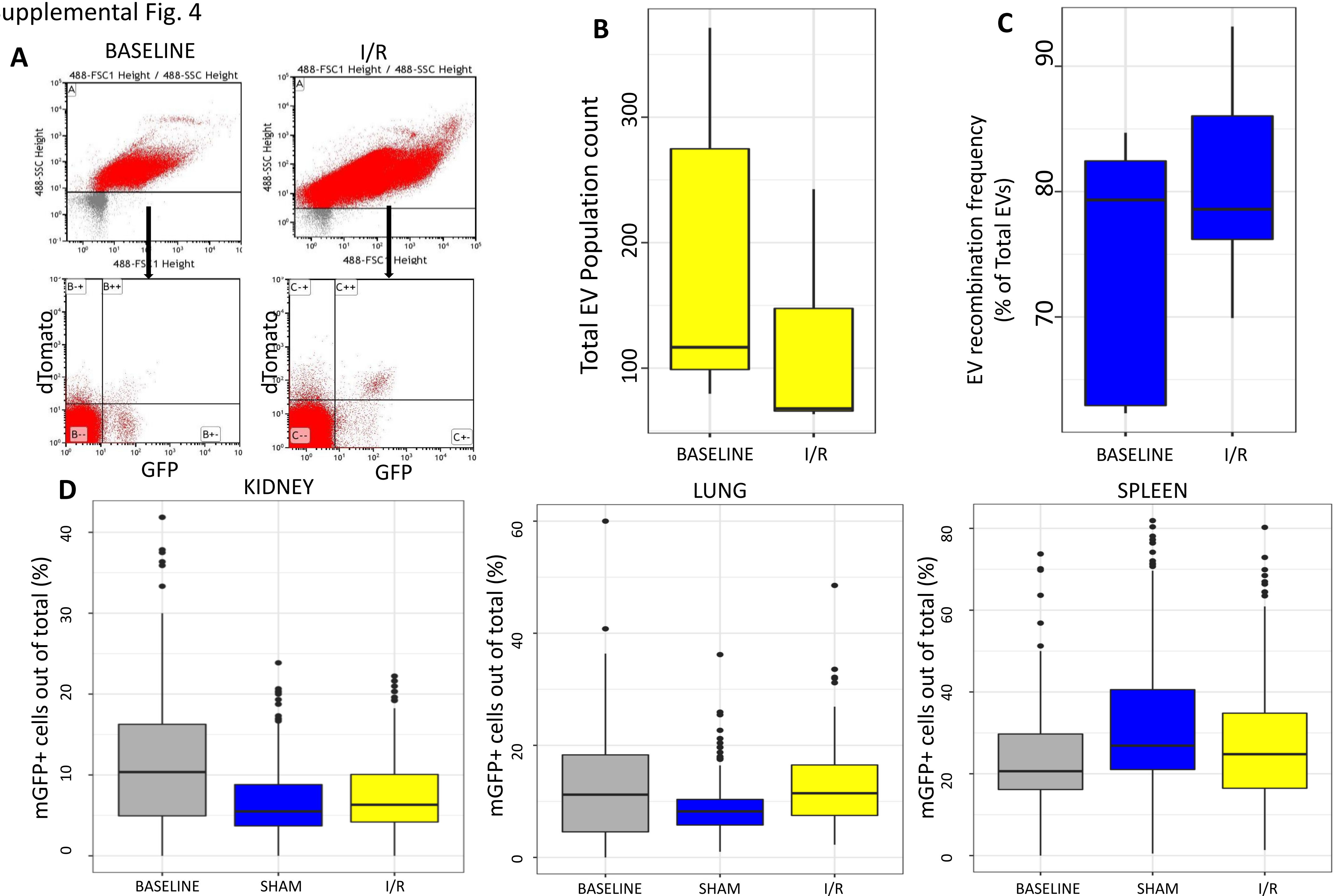

Supplemental Fig. 5

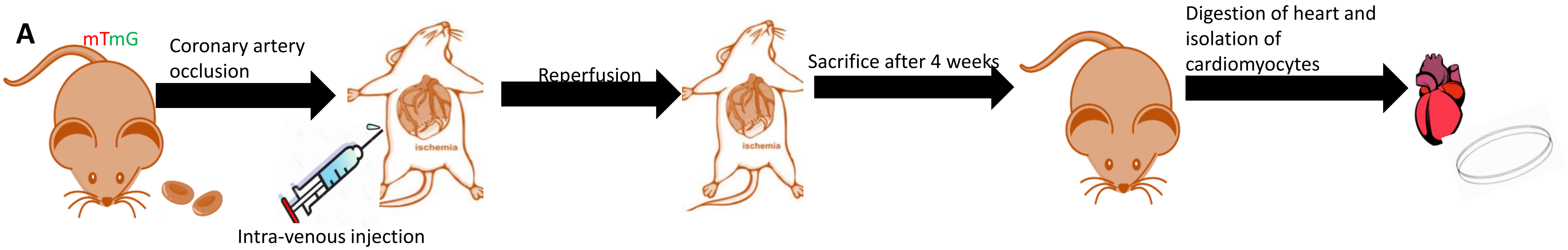


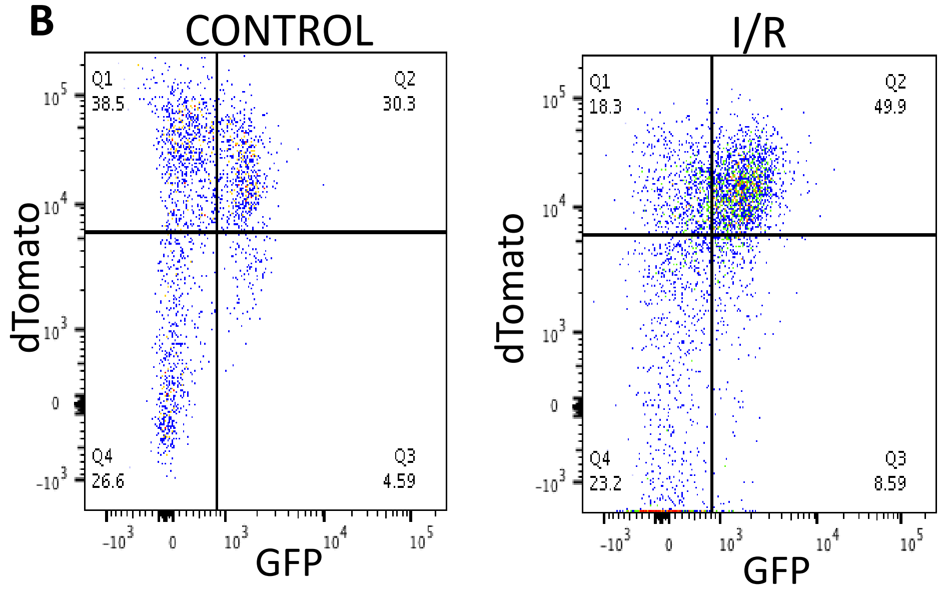


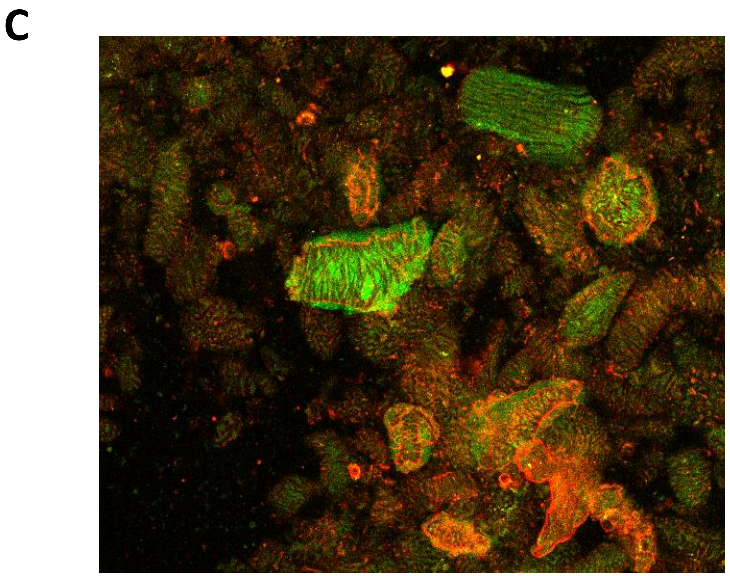

Supplemental Fig. 6

**A**

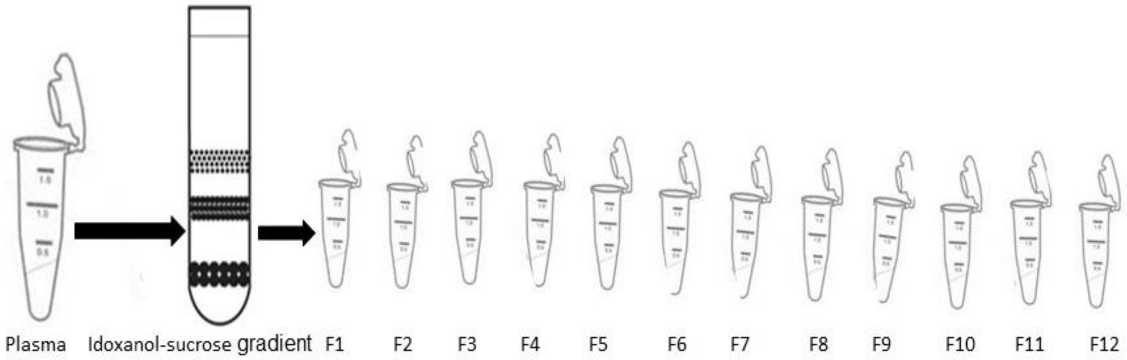


**B**

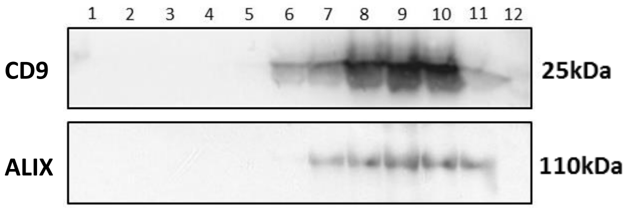


**C**

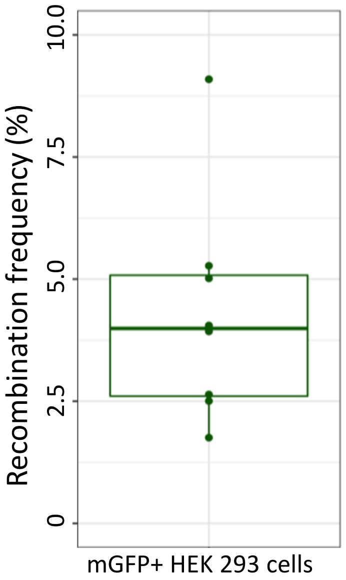

Supplemental Fig. 7

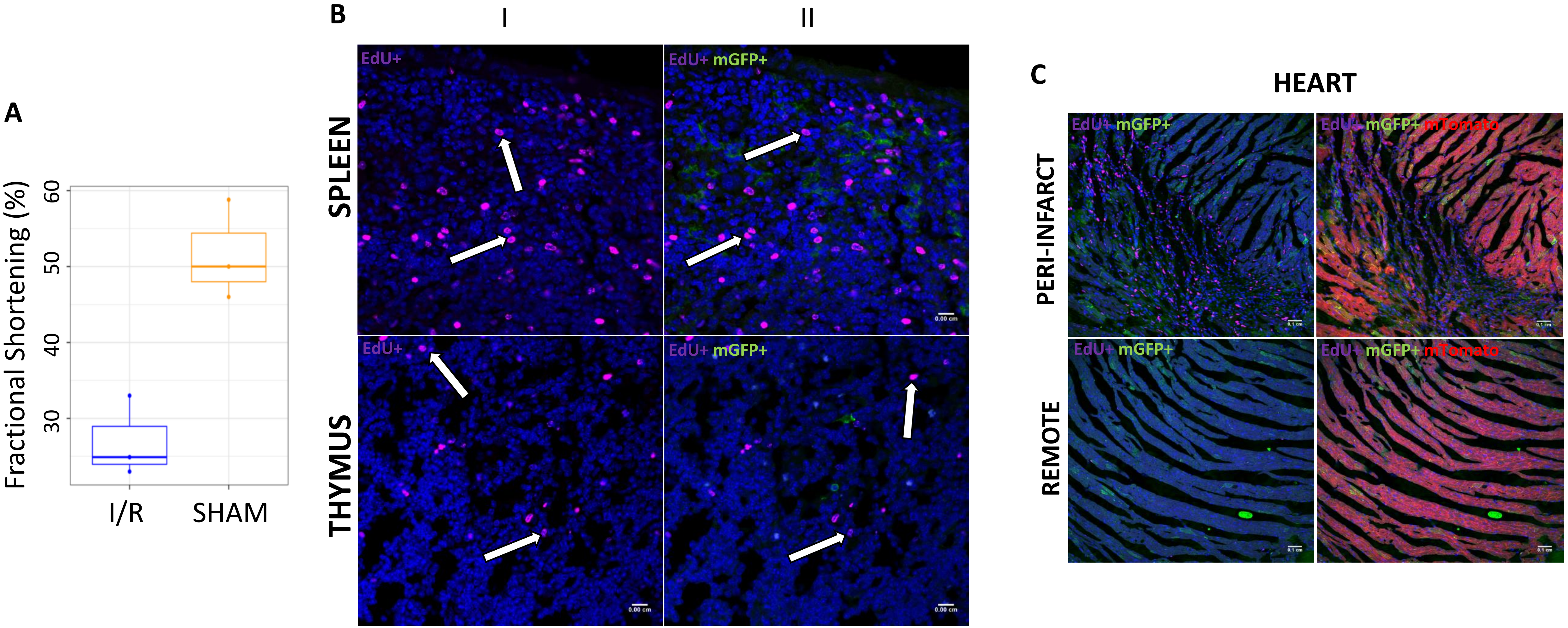
